# Nimbolide Targets RNF114 to Induce the Trapping of PARP1 and Poly-ADP-Ribosylation-Dependent DNA Repair Factors

**DOI:** 10.1101/2022.10.04.510815

**Authors:** Peng Li, Yuanli Zhen, Chiho Kim, Heping Deng, Hejun Deng, Xu-Dong Wang, Tian Qin, Yonghao Yu

## Abstract

PARP1 is an abundant nuclear enzyme that is critically involved in DNA damage response. Its main enzymatic function is to catalyze a protein post-translational modification known as poly-ADP-ribosylation (PARylation). Despite the tremendous progresses of PARP1 inhibitors (PARPi) in the clinic, the basic mechanism of action of PARPi is poorly understood. Recent studies point to PARP1 trapping as a key factor driving the cytotoxic and immunomodulatory functions of PARPi. However, the molecular underpinnings of PARP1 trapping remain elusive. Here, using an unbiased, quantitative proteomic screen, we identified RING finger protein 114 (RNF114), as a PARylation-dependent, ubiquitin E3 ligase involved in DNA damage response. Upon sensing DNA damage, RNF114 was recruited, in a PAR-dependent manner, to the DNA lesions, where it specifically targeted PARylated-PARP1 for ubiquitin-proteasomal degradation. The blockade of this pathway interfered with the removal of PARP1 from the DNA damage site, leading to profound PARP1 trapping. We show that a natural product, Nimbolide, targeted RNF114 to block the degradation and removal of PARylated-PARP1 from DNA lesions. Unlike conventional PARPi, nimbolide treatment induced the trapping of both PARP1 and PARylation-dependent DNA repair factors. This unique mechanism of action of nimbolide translated into its superior cytotoxic effects against *BRCA*-deficient cancers. Furthermore, we demonstrated that, as a super trapper of both PARylated-PARP1 and PARylation-dependent DNA repair factors, nimbolide was able to overcome the intrinsic and acquired resistance to PARPi. Finally, we showed that nimbolide treatment activated innate immune signaling and was able to synergize with various cytotoxic agents. These results point to the exciting possibility of targeting homologous recombination-deficient cancers using nimbolide and its analogs.

## Introduction

Poly-ADP-ribosylation (PARylation) is a protein posttranslational modification (PTM) that was first documented in 1963^1^. PAR is synthesized by a class of enzymes called poly-ADP-ribose polymerases (PARPs), which utilize NAD^+^ as a cofactor. Among the various PARP enzymes, PARP1 is an abundant nuclear enzyme that functions as a critical DNA damage sensor^2–5^. In response to genotoxic stress, PARP1 is recruited to nicked DNA^6^. The binding of PARP1 to a DNA strand break induces a profound conformational change of PARP1, resulting in the remodeling of an inhibitory motif near the catalytic domain, and hence stimulation of its enzymatic activity^7^. The activated PARP1 then catalyzes the PARylation on a large array of substrate proteins (including PARP1 itself), leading to the initiation of DNA damage response (DDR)^8^.

PAR polymers are composed of linear and/or branched repeating units of ADP-ribose, and these PAR chains are bulky, charged and flexible^3^. As a result, PARylation could regulate the function of a target protein by modulating its electrostatic and topological properties^9,10^. Furthermore, protein-linked PAR polymers may also serve as a platform for the recruitment of other signaling molecules. Indeed, a number of PAR-binding motifs (PBMs) (e.g., the PBZ, BRCT, WWE, FHA, OB-fold, RRM and macro domain) have been identified, and these PBMs are present in many proteins involved in DDR^2,4^. Among them, DNA damage-induced PAR chains recruit XRCC1 via the binding of its BRCT domain to these PAR polymers. XRCC1 then recruits additional protein factors (e.g., DNA ligase IIIα/LIG3 and DNA Polβ), thereby seeding the formation of a large protein complex that mediates the repair of DNA single-strand breaks (SSBs)^11,12^.

*BRCA1/2*-mutated cancer cells are homologous recombination-deficient, and it has been shown that they are selectively killed by PARP1 inhibitor (PARPi), owing to the synthetic lethality mechanism^13,14^. To date, four PARPi (Olaparib, Rucaparib, Niraparib, and Talazoparib) have been approved by the FDA for the treatment of breast, ovarian, pancreatic and/or prostate cancers with homologous recombination-deficiency (HRD)^15^. Despite these tremendous progresses of PARPi in the clinic, the exact mechanism of action (MoA) of PARPi remains poorly understood. Furthermore, intrinsic and acquired resistance to PARPi is frequently observed in the clinic, which points to the critical need for the development of next generation PARP1-targeting agents with more complete and sustained response.

All FDA-approved PARPi competitively occupy the NAD^+^-binding pocket of PARP1. These compounds, however, simultaneously possess two distinct, yet interconnected activities (i.e., PARP1 inhibition and PARP1 trapping)^16,17^. By inhibiting the catalytic activity of PARP1, PARPi could kill tumor cells by blocking PARylation-dependent DDR signaling^15^. However, PARPi with similar PARP1 inhibitory activities have dramatically different cytotoxicity^18,19^. The MoA of PARPi, therefore, has been reconsidered beyond the initial hypothesis that PARPi acts as DNA repair inhibitors by blocking the catalytic activity of PARP1. More recent studies then showed that besides inhibiting PARP1, all FDA-approved PARPi also induce PARP1 trapping^18^. The recruitment of PARP1 to DNA lesions represents one of the earliest molecular events during DDR^2,20^. Upon the treatment of PARPi, PARP1, however, is retained at the DNA damage site for an extended time (termed “trapping”), resulting in the formation of a trapped PARP1-DNA-PARPi complex^6,18^. The trapped PARP1 causes the collapse of the replication fork and is known to be highly toxic to cells^6^. Results from recent genetic and pharmacological experiments showed that the presence of the PARP1 protein with uncompromised DNA-binding activities is required for PARPi-induced DDR, cytotoxicity, and innate immune signaling^18,21–24^. Furthermore, it has been shown that the cytotoxicity of various PARPi is positively correlated with their ability to induce PARP1 trapping*^25^*. These results suggest that PARP1 trapping might function as the key determinant for the anti-tumor effects of PARPi^17^. However, at the molecular level, PARP1 inhibition and PARP1 trapping are coupled (e.g., via PARP1 auto-PARylation), and the mechanistic nature of PARP1 trapping remains poorly understood.

Here we performed an unbiased, quantitative mass spectrometry-based screen to identify protein factors that are re-localized, in a PARylation-dependent manner, to the chromatin during DDR. Through these experiments, we garnered unanticipated insights into the molecular underpinnings of PARP1 trapping. Specifically, from the proteomic screen, we identified a poorly studied ubiquitin E3 ligase, RNF114, that showed PARylation-dependent recruitment to DNA lesions. We further showed that PAR chains not only bind to RNF114, but also stimulate the E3 ligase activity of this enzyme. Using a series of biochemical assays, we identified PARP1 as a novel substrate of RNF114, and importantly, RNF114 targets PARylated-PARP1 for ubiquitination and proteasomal degradation. Genetic deletion of RNF114, or inhibition of its E3 activity leads to potent PARP1 trapping. These results suggest that blockade of the ubiquitination and proteasomal degradation of PARP1 could be a key contributing factor of PARP1 trapping. Nimbolide, a natural product derived from the Neem tree (*Azadirachta indica*), targets the ubiquitin E3 ligase activity of RNF114, and thereby induces the trapping of PARP1. Although regular NAD^+^-competitive PARPi induce the trapping of PARP1, nimbolide treatment causes the trapping of PARylated-PARP1, and also PAR-dependent DNA repair factors. We showed that this unique mechanism of action of nimbolide translates into its enhanced cytotoxicity in *BRCA*-deficient tumor cells, and also its ability to kill tumor cells with intrinsic and acquired resistance to conventional PARPi. We found that nimbolide synergizes with various DNA damaging agents. Finally, by inducing PARP1 trapping and DDR, nimbolide activates innate immune signaling. These results point to a model where RNF114-mediated ubiquitination and degradation of PARP1 is involved in the removal of PARP1 from DNA lesions. By blocking the RNF114-dependent PARP1 degradation pathway, nimbolide treatment induces the trapping of both PARP1 and PARylation-dependent DNA repair factors. This unique mechanism of action points to the exciting possibility of translating nimbolide and its analogs into novel therapeutic agents against *BRCA*-deficient cancers.

## Results

### Comprehensive Identification of the Poly-ADP-Ribosylation (PARylation)-Dependent DNA Repair Factors

We performed a chromatin relocation screen to identify the factors involved in the PARylation-dependent DNA damage response (Fig. 1a). In this case, we used quantitative mass spectrometric profiling experiments to comprehensively characterize the chromatin-associated proteome in cells treated with genotoxic agents that activate PARP1. We pre-treated the HCT116 cells with DMSO or Talazoparib (a potent PARPi)^26^. The cells were then treated with H_2_O_2_ or MMS to induce DNA damage, and the subsequent PARP1 activation. Immunoblot analyses showed that the treatment of H_2_O_2_ or MMS induced potent PARP1 activation, which, however, was blocked by Talazoparib pre-treatment (Fig. 1b). For the quantitative proteomic profiling experiments, we harvested the treated cells and isolated the chromatin fraction. After protein extraction and proteolytic digestion, we used isobaric labeling-based quantitative mass spectrometry for global protein expression profiling experiments^27^. Specifically, the digested peptides were labeled with the corresponding TMT (tandem mass tag) reagent. These samples were combined, which were then subject to multidimensional HPLC separation coupled with quantitative mass spectrometric experiments^28^. From the combined dataset for the quantitative proteomic MS experiments, we were able to identify and quantify a total of 2,346 proteins from these chromatin fractions (Supplementary Table 1).

**Fig. 1.**
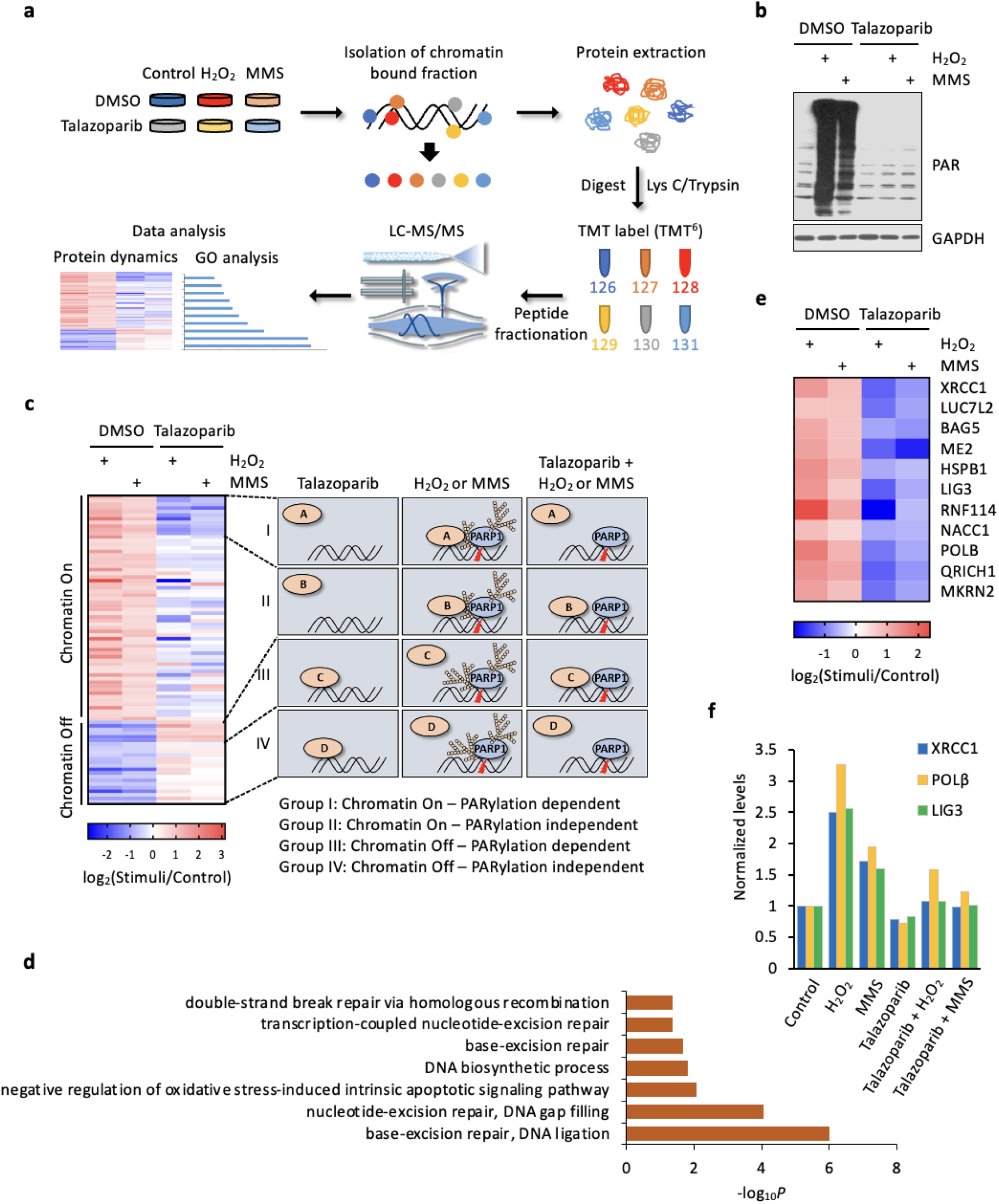
A chromatin re-localization screen for the identification of Poly-ADP-ribosylation (PARylation)-dependent DNA repair factors. **a,** Overall scheme of the experimental procedures for chromatin re-localization screen. **b,** Immunoblot analyses of the whole cell lysate samples derived from (**a**). **c,** Hierarchical clustering of the re-localized proteins in response to genotoxic stress. The identified proteins were further classified into four categories, i.e., group I (Chromatin on; PARylation dependent); group II (Chromatin on; PARylation independent); group III (Chromatin off; PARylation dependent); and group IV (Chromatin off; PARylation independent). See the main text for the detailed description of these four classes of proteins. **d,** GO analyses of the group I proteins (Chromatin-On; PARylation-dependent proteins) identified from the chromatin re-localization screen. **e,** A detailed heat map of all the group I proteins. **f,** The abundances of representative group I proteins (i.e., XRCC1, POLβ, and LIG3) in the chromatin fraction as determined from the chromatin re-localization screen.

The identified proteins were subject to unsupervised hierarchical clustering analyses (Fig. 1c, Extended Data Fig. 1). The results showed that these proteins converged into four clusters: Group I: Chromatin-On, PARylation-dependent proteins, i.e., those with increased abundances (log_2_FC > 1; FC, fold-of-change) in the chromatin samples isolated from the MMS/H_2_O_2_-treated cells, compared to control cells. The increase, however, was abolished in the cells that were pre-treated with Talazoparib; Group II: Chromatin-On, PARylation-independent proteins, i.e., those with increased abundances (log_2_FC > 1) in the chromatin samples isolated from the MMS/H_2_O_2_-treated cells, compared to control cells. The increase, however, was not affected by Talazoparib pre-treatment; Group III: Chromatin-Off, PARylation-dependent proteins, i.e., those with decreased abundances (log_2_FC < −1) in the chromatin samples isolated from MMS/H_2_O_2_-treated cells, compared to control cells. The decrease, however, was abolished in the cells that were pre-treated with Talazoparib; and Group IV: Chromatin-Off, PARylation-independent proteins, i.e., those with decreased abundances (log_2_FC < −1) in the chromatin samples isolated from MMS/H_2_O_2_-treated cells, compared to control cells. The decrease, however, was not affected by Talazoparib pre-treatment.

We performed biochemical experiments to validate the results from our quantitative proteomic screen. We selected several representative hits from our screen (e.g., INTS5, INTS3, SOX11, ZC3H3 and TAF10), and the results showed that an excellent agreement was achieved between the proteomic and biochemical experiments (Extended Data Fig. 2). In summary, our chromatin proteomic screen serves as an invaluable approach for the identification of potential PARylation-dependent and -independent DNA repair factors.

### Identification of RNF114 as a PAR-binding ubiquitin E3 ligase

Among the four clusters, we were focused on the Group I proteins (i.e., the chromatin-on, PARylation-dependent proteins) (Fig. 1e). We performed gene ontology (GO) analyses of this group of proteins (Fig. 1d, Extended Data Fig. 1) and found that most of these proteins were nuclear proteins involved in DDR-related biological processes, including base-excision repair and DNA ligation (*P* = 9.57 × 10 ^−7^), and nucleotide-excision repair and DNA gap filling (*P* = 8.75 × 10^−5^) (Fig. 1d). This group contained several DNA repair factors (e.g., XRCC1, POLβ and LIG3) that are known to be recruited, in a PARylation-dependent manner, to chromatin in response to DNA damage (Fig. 1e, f). The identification of these known PARylation-dependent DNA repair factors demonstrated the validity of our screen.

Among the other Group I proteins, we were particularly intrigued by a protein called RNF114. RNF114 (also known as ZNF313) is a poorly studied E3 ubiquitin-protein ligase with largely unknown functions^29,30^. A previous study showed that RNF114 interacts with A20 and modulates the NF-γB pathway and T-cell activation^31^. However, the role of RNF114 in DDR has not been defined. We performed independent biochemical studies and showed that RNF114 was indeed recruited to the chromatin during DDR (Fig. 2a). Its recruitment, however, was blocked in cells that were pre-treated with PARPi. Our proteomic and biochemical studies, therefore, suggest that RNF114 could be a potential factor involved in PARylation-dependent DNA damage response.

**Fig. 2.**
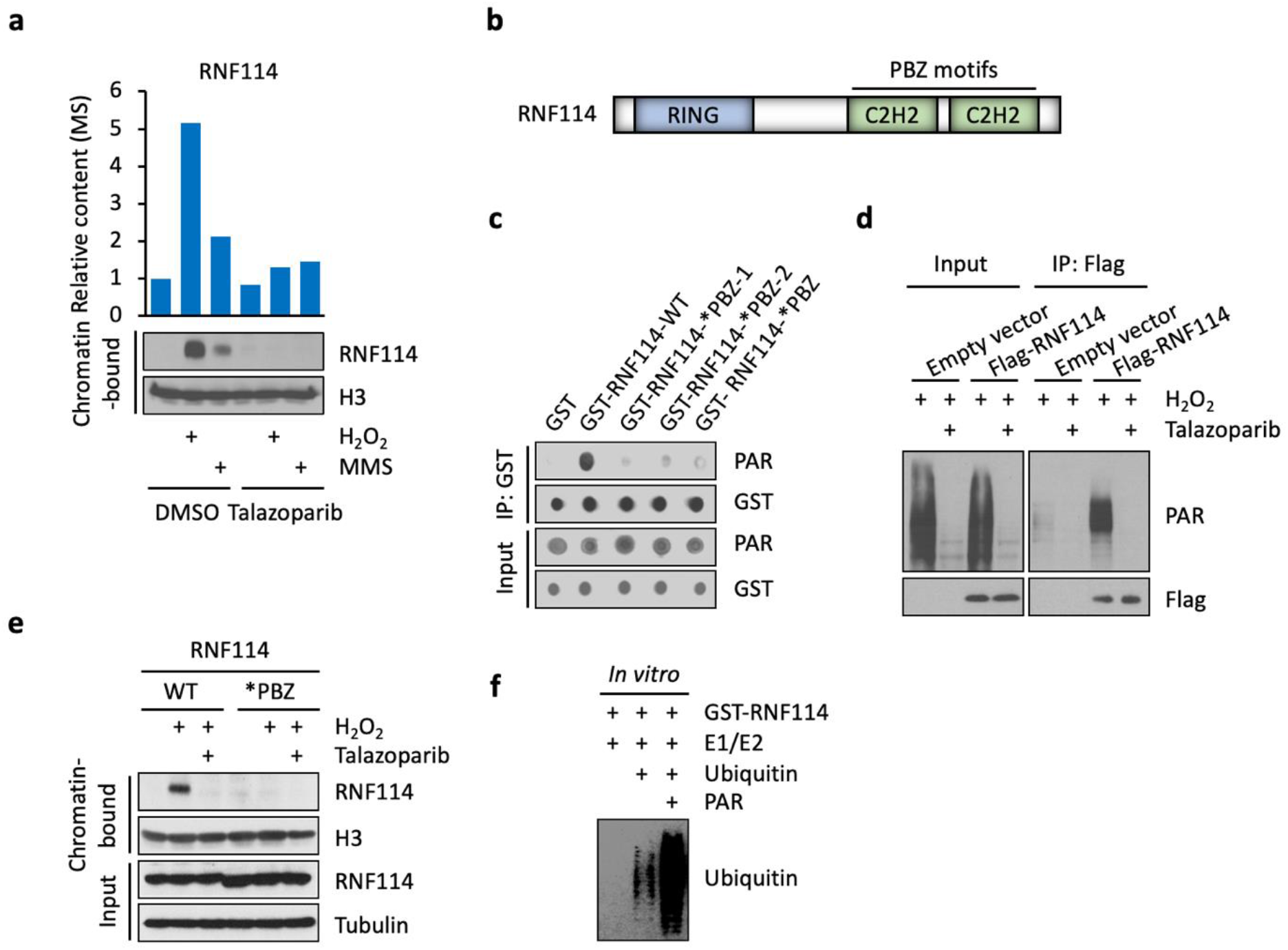
Identification of RNF114 as a PAR-binding, ubiquitin E3 ligase. **a,** The abundances of RNF114 in the chromatin fraction as determined from the chromatin re-localization screen (upper panel) and biochemical assays (lower panel). **b,** The domain structure of RNF114. **c, d,** RNF114 binds to PAR polymers *in vitro* (**c**) and in cells (**d**). **c,** Both RNF114-WT and the various RNF114 mutants were analyzed using the PAR dot-blot assay (i.e., RNF114-WT; RNF114-*PBZ-1, C143A/C146A; RNF114-*PBZ-2, C173A/C176A; RNF114-*PBZ, C143A/C146A/C173A/C176A). **d,** HCT116 cells expressing the empty vector or Flag-RNF114 were pre-treated with Talazoparib (1 μM for 1 h), and then were treated with H_2_O_2_ (2 mM for 5 min). Cell lysates were subject to immunoprecipitation using an anti-FLAG antibody, and the immunoprecipitants were analyzed using the indicated antibodies. **e,** PAR binding mediates the translocation of RNF114 to chromatin in response to genotoxic stress. RNF114-KO HCT116 cells were reconstituted with either RNF114-WT or RNF114-*PBZ mutant (the PAR-binding mutant). These cells were pre-treated with Talazoparib (1 μM for 1 h), and then were treated with H_2_O_2_ (2 mM for 5 min). The chromatin-bound fraction was isolated from these cells, and was subject to immunoblotting experiments using the indicated antibodies. **f,** PAR polymers stimulate the E3 activity of RNF114. Recombinant RNF114 was subject to *in vitro* ubiquitination experiments in the presence (0.4 μM) or absence of added PAR polymers. Immunoblot experiments were used to analyze RNF114 auto-ubiquitination.

RNF114 has several distinct protein domains, including an amino-terminal RING domain (an E3 ligase domain), which is followed by two C2H2 [Cys(2)-His(2)]-type zinc finger domains (Fig. 2b). We then used a dot-blot assay to biochemically test whether RNF114 is a PAR-binding protein. We found that only RNF114 wild-type (WT), but not the RNF114 PBZ mutants (*PBZ-1, C143A/C146A; *PBZ-2, C173A/C176A; *PBZ, C143A/C146A/C173A/C176A) interacted with PAR polymers (Fig. 2c). Furthermore, we treated cells with H_2_O_2_ to induce DNA damage and to initiate PARylation. Consistent with the notion that RNF114 is a potential PAR-binding protein, a strong PAR signal was detected in the RNF114 immunoprecipitants (Fig. 2d). Furthermore, we performed biochemical fractionation experiments and found that either pre-treatment of Talazoparib or PBZ mutation completely abolished H_2_O_2_-induced chromatin translocation of RNF114 (Fig. 2e). Together, these findings suggest that the recruitment of RNF114 to the DNA lesions is dependent on its PAR binding domains.

Previous studies reported that RNF114 is an E3 ligase that is able to modify itself and certain substrate proteins (e.g., A20 and p21) by ubiquitination^29,32,33^. Consistent with these studies, we found that RNF114 was able to catalyze its auto-ubiquitination in an *in vitro* ubiquitination assay (Fig. 2f). Because RNF114 interacts with PAR chains, we further tested whether PAR is a regulator of the E3 ligase activity of RNF114. We found that PAR chains dramatically stimulated the E3 ligase activity of RNF114 (Fig. 2f), suggesting that RNF114 is a PAR-dependent E3 ligase.

### RNF114 targets PARylated-PARP1 for ubiquitin-proteasomal degradation

Although RNF114 is known to ubiquitinate several proteins (e.g., A20 and p21)^29,31^, its substrate profile is poorly defined. In particular, it is unclear whether RNF114 targets certain proteins for ubiquitination and degradation, in order to mediate its potential functions in DDR. To identify the substrates of RNF114 in the context of PARylation-mediated DDR, we performed an IP-MS (immunoprecipitation coupled with MS) experiment, using cells treated with either H_2_O_2_ or H_2_O_2_+Talazoparib. We identified a large number of PARP1 peptides from the RNF114 immunoprecipitants only in H_2_O_2_-treated cells (Fig. 3a). These results were subsequently validated using immunoblotting experiments (Fig. 3a). However, the interaction between RNF114 and PARP1 was completely blocked by Talazoparib pre-treatment (Fig. 3a). Consistent with the notion that RNF114 is a PAR-binding protein, these results indicate that RNF114 binds to PARylated-PARP1, but not PARP1.

**Fig. 3.**
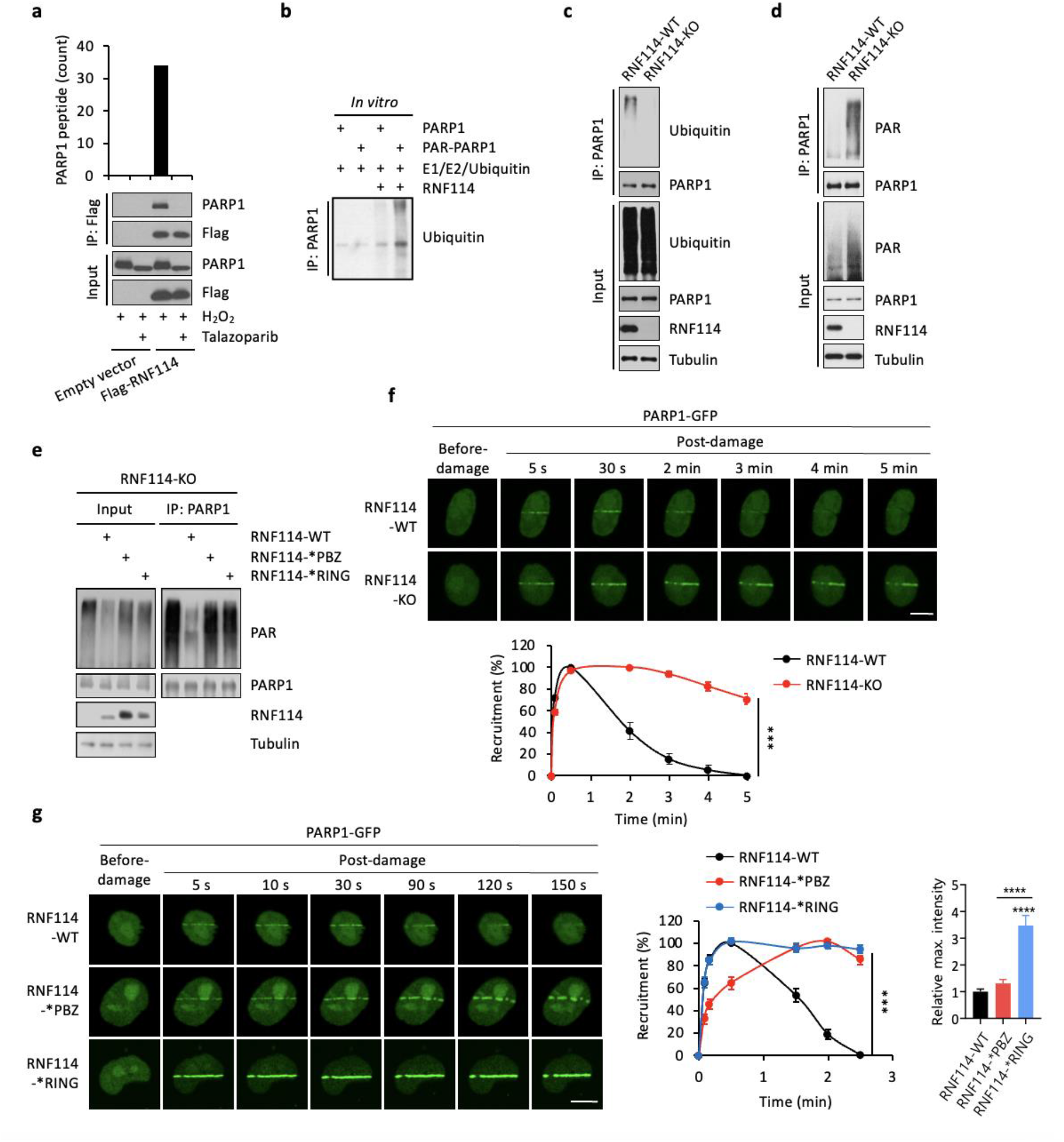
RNF114 targets PARylated-PARP1 for ubiquitin-proteasomal degradation. **a,** RNF114 interacts with PARylated-PARP1. HCT116 cells expressing the empty vector or Flag-RNF114 were pre-treated with Talazoparib (1 μM for 1 h), and were treated with H_2_O_2_ (2 mM for 5 min). The whole cell lysates were subject to immunoprecipitation (anti-Flag), and the immunoprecipitants were analyzed by LC-MS/MS experiments (upper panel) and immunoblot experiments (lower panel). **b,** *In vitro* ubiquitination assays of PARP1 or PARylated-PARP1. The assay was performed using the protocol as shown in Fig. 2f, with PARP1 or PARylated PARP1 as the substrate (both were used at 1 μg). After the in vitro ubiquitination reaction, the samples were denatured by the addition of 1% SDS (final concentration) and were boiled. The samples were diluted (10X) using the lysis buffer (to reduce the concentration of SDS to 0.1%), and were subject to immunoprecipitation using the PARP1 antibody (to remove the interference from RNF114). The isolated PARP1 was probed using the anti-ubiquitin antibody. **c,** RNF114 mediates the ubiquitination of PARP1. Control (RNF114-WT) and RNF114-KO HCT116 cells were pre-treated with MG132 (10 μM for 6 h), and then were treated with H_2_O_2_ (2 mM for 5 min). PARP1 was isolated using immunoprecipitation, and was subject to immunoblotting analyses using the indicated antibodies. **d,** RNF114 mediates the degradation of PARylated-PARP1. Control (RNF114-WT) and RNF114-KO HCT116 cells were treated with H_2_O_2_ (2 mM for 5 min). PARP1 was isolated using immunoprecipitation, and was subject to immunoblotting analyses using the indicated antibodies. **e,** RNF114 with the uncompromised PAR-binding and E3 ligase activity is required for the degradation of PARylated-PARP1. RNF114-KO HCT116 cells were reconstituted with RNF114-WT, RNF114-*PBZ mutant, or RNF114-*RING mutant. These cells were pre-treated with H_2_O_2_ (2 mM for 5 min). PARP1 was isolated using immunoprecipitation, and was subject to immunoblotting analyses using the indicated antibodies. **f,** Deletion of RNF114 leads to PARP1 trapping. Control (RNF114-WT) and RNF114-KO HeLa cells expressing GFP-tagged PARP1 were subject to the laser microirradiation experiment. GFP signals were monitored in a time-course experiment with the quantification shown (lower panel). Scale bars, 10 μm. **g,** Interference with the PAR-binding or the E3 activity of RNF114 leads to PARP1 trapping. RNF114-KO HeLa cells were reconstituted with the RNF114-WT, RNF114-*PBZ mutant, or RNF114-*RING mutant. These cells were transfected with GFP-tagged PARP1, and were subject to a laser microirradiation assay. GFP signals were monitored and were quantified in a time-course experiment (right panel). Scale bars, 10 μm. The level of PARP1 trapping (relative max intensity) was quantified by dividing the GFP signal from trapped PARP1 (trapped PARP1-GFP, combined across the different time points) by the total GFP signal in the nucleus (total PARP1-GFP, combined across the different time points). The level of PARP1 trapping for the different RNF114 conditions was then normalized to that in the RNF114-WT sample.

Next, we tested the hypothesis that RNF114 could target PARylated-PARP1 for ubiquitin and its subsequent proteasomal degradation. We performed *in vitro* ubiquitination experiments and found that RNF114 ubiquitinated the PARylated-PARP1 (PAR-PARP1) (Fig. 3b). To further investigate this in intact cells, we pre-treated control or RNF114-KO cells with MG132 to block protein degradation. We then stimulated these cells with H_2_O_2_ to activate PARP1, and isolated PARP1 using immunoprecipitation. We found that the ubiquitination signal in PARP1 immunoprecipitants was dramatically decreased in RNF114-KO cells, compared to that in the control cells (Fig. 3c). These results suggest that RNF114 is a major E3 ligase that mediates the ubiquitination of PARP1 under genotoxic conditions. In another experiment, we treated control or RNF114-KO cells with H_2_O_2_ to induce DNA damage and the activation of PARP1. Consistent with the potential role of RNF114 in mediating the degradation of PARylated PARP1, we found that the PAR signal in PARP1 immunoprecipitants was dramatically decreased in control cells, compared to that in the RNF114-KO cells (Fig. 3d).

Since the PBZ domain is a PAR-binding motif (Fig. 2c)^4^, we first confirmed that PARylated-PARP1 was bound strongly to RNF114-WT or the RNF114-*RING mutant, but not the RNF114-*PBZ mutant (Extended Data Fig. 3a). Next we reconstituted the RNF114-KO cells with RNF114-WT, RNF114-*RING (the RING mutant, C29A/C32A, compromised in its E3 ligase activity)^34^, or RNF114-*PBZ (the PBZ mutant, C143A/C146A/C173A/C176A, compromised in its PAR-binding ability). PARylated-PARP1 was only degraded in cells expressing RNF114-WT, but not those expressing the RNF114-*RING mutant, or the RNF114-*PBZ mutant (Fig. 3e). The degradation of PARylated-PARP1 was completely blocked by MG132 pre-treatment, suggesting that PARylated-PARP1 was degraded via the ubiquitin-proteasomal system (Extended Data Fig. 3b). Taken together, these results demonstrate that RNF114 is a PAR-dependent E3 ligase that targets PARylated-PARP1 for degradation via the ubiquitin proteasome pathway.

### RNF114 blockade results in PARP1 trapping

Using the laser microirradiation assays, we found that RNF114 co-localized with PCNA upon micro-irradiation, further suggesting that RNF114 is involved in the DNA damage repair process (Extended Data Fig. 3c). Next, we performed laser microirradiation assays to examine how RNF114 regulates the recruitment and retention of PARP1 during DDR. In control cells, PARP1 was recruited to DNA lesions within a few seconds and then the PARP1 signal disappeared from DNA lesions after ~5 min (Fig. 3f). However, PARP1 remained at the DNA lesions for a prolonged time in RNF114-KO cells (Fig. 3f), which is consistent with the formation of trapped PARP1. These data are also consistent with a model where RNF114 is recruited, in a PARylation-dependent manner, to the DNA damage site, where it removes PARP1 via the ubiquitin-proteasomal mechanism. The blockage of this pathway instead causes PARP1 trapping.

Additionally, we also examined the kinetics of PARP1 relocation in the RNF114-KO cells reconstituted with RNF114-WT, the RNF114-*PBZ mutant or the RNF114-*RING mutant. Compared to RNF114-WT cells, PARP1 in RNF114-*PBZ mutant cells was retained on DNA lesions for a prolonged time (i.e., PARP1 trapping). This is likely due to the PAR binding deficiency and hence the compromised PAR-mediated recruitment of this RNF114 mutant (Fig. 3g). PARP1 was also retained at the DNA lesions for a prolonged time in cells expressing the RNF1114-*RING mutant. This is likely due to its compromised E3 ligase activity (Fig. 3g). The RNF114-*RING mutant induced a stronger level of PARP1 trapping, compared to the RNF114-*PBZ mutant (Fig. 3g). These data are consistent with a potential dominant-negative effect of the RNF114-*RING mutant on PARP1 trapping. Specifically, it is likely that even though this mutant does not degrade PARylated-PARP1, it occupies PARylated-PARP1 through the intact PBZ motifs. This binding could protect PARylated-PARP1, and prevent it from being removed from DNA lesions by other PAR-dependent mechanisms.

PARP1 trapping causes the stalling and collapse of replication forks, and is known to be the main mechanism driving the cytotoxicity of PARPi^18^. Consistent with the role of RNF114 in regulating PARP1 trapping, we found that compared to control cells, RNF114-KO cells were more susceptible to various genotoxic agents (e.g., H_2_O_2_) (Extended Data Fig. 3d). Furthermore, MMS-induced cell death was ameliorated in RNF114-KO cells reconstituted with RNF114-WT, but not the RNF114-*RING mutant or the RNF114-*PBZ mutant (Extended Data Fig. 3e). Consistent with the ability of RNF114-*RING to induce potent PARP1 trapping, cells expressing the RNF114-*RING mutant showed the most profound levels of cell death under genotoxic conditions (Extended Data Fig. 3e).

### Nimbolide traps PARP1 and the PAR-dependent DNA repair factors

Nimbolide is a natural product that was originally isolated from the Neem tree (*Azadirachta indica*) (Fig. 4a)^35^. Previous studies suggest that nimbolide has certain anti-cancer activities, although its underlying mechanism of action is poorly understood^36–39^. A recent study showed that nimbolide covalently modifies RNF114. It has been suggested that this binding event blocks the substrate engagement of RNF114, and therefore prevents the ubiquitination and the subsequent degradation of its target substrates^40^. Consistent with this model, we found that nimbolide was able to potently block the auto-ubiquitination of RNF114 in an in vitro ubiquitination assay (Fig. 4f, Extended Data Fig. 3f).

**Fig. 4.**
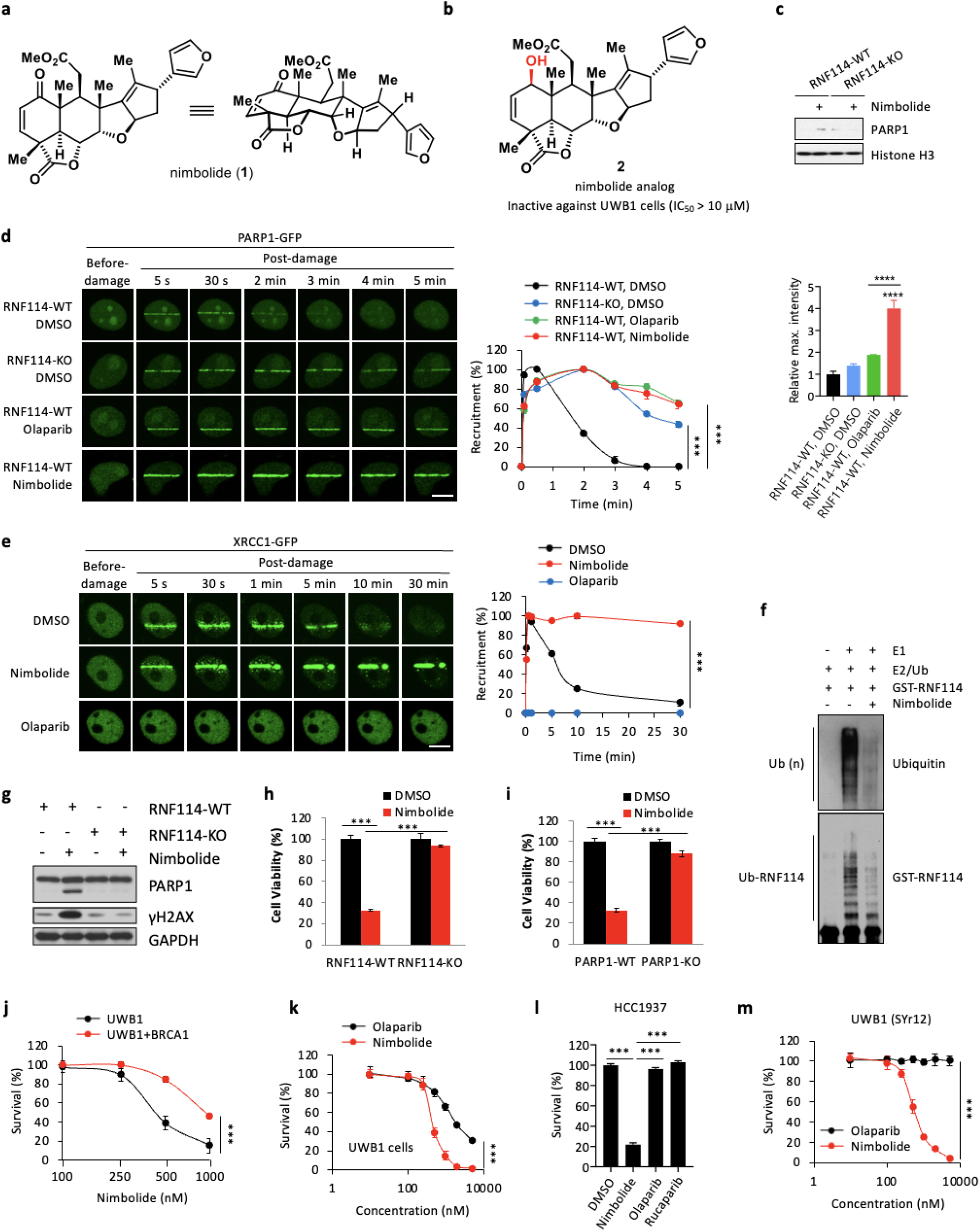
Nimbolide traps PARylated-PARP1 and PAR-dependent DNA repair factors. Structure of nimbolide (**a**) and a nimbolide analog (**b**). The compound as shown in (**b**) was inactive against UWB1 cells (IC_50_ > 10 μM). Additional nimbolide analogs are shown in Extended Data Fig. 4c. **c,** RNF114 deletion abolishes the nimbolide-induced PARP1 trapping. HCT116 control (RNF114-WT) or HCF116 RNF114-KO cells were treated with nimbolide (1 μM for 24 hrs). The cells were subject to subcellular fractionation, and the chromatin fraction was analyzed by immunoblot assays using the indicated antibodies. **d,** Nimbolide treatment induces potent PARP1 trapping. Control (RNF114-WT) or RNF114-KO HeLa cells expressing GFP-tagged PARP1 were pre-treated with Olaparib (10 μM) or nimbolide (1 μM) for 1 h. GFP signals were monitored in a time-course experiment following laser microirradiation. The quantification results are shown in the right panel. Scale Bars, 10 μm. The level of PARP1 trapping (relative max intensity) was quantified by dividing the GFP signal from trapped PARP1 (trapped PARP1-GFP, combined across the different time points) by the total GFP signal in the nucleus (total PARP1-GFP, combined across the different time points). The level of PARP1 trapping for the different RNF114 conditions was then normalized to that in the RNF114-WT, DMSO sample. **e,** Nimbolide, but not Olaparib treatment, induces the trapping of XRCC1. HeLa cells expressing GFP-tagged XRCC1 were pre-treated with Olaparib (1 μM) or nimbolide (1 μM) for 1 h. These cells were subject to a laser microirradiation assay. GFP signals were monitored and were calculated in a time course. The quantification results are shown in the right panel. Scale bars, 10 μm. **f,** Nimbolide blocks the auto-ubiquitination of RNF114 in an in vitro ubiquitination assay. Where indicated, GST-RNF114 was incubated with E1/E2/ubiquitin in the presence of nimbolide (1 μM). **g,** RNF114 deletion rescues cells from nimbolide-induced cell death. Control (RNF114-WT) or RNF114-KO HeLa cells were treated with nimbolide (1 μM) for 24 hrs. Cells were lysed and the whole cell lysates were subject to immunoblotting analyses using the indicated antibodies. The deletion of RNF114 was confirmed as shown in Extended Data Fig. 3g. **h,** Control (RNF114-WT) or RNF114-KO HeLa cells were treated with nimbolide (1 μM) for 24 hrs, and cell viability was quantified using the CellTiter-Glo assay. The deletion of RNF114 was confirmed as shown in Extended Data Fig. 3g. **i,** PARP1 deletion rescues cells from nimbolide-induced cell death. Control (PARP1-WT) or PARP1-KO HeLa cells were treated with nimbolide (1 μM) for 24 hrs, and cell viability was quantified using the CellTiter-Glo assay. The deletion of PARP1 was confirmed as shown in Extended Data Fig. 3g. **j,** Nimbolide treatment is synthetic lethal with respect to *BRCA1*-deficiency. UWB1 and UWB1+BRCA1 cells were subject to nimbolide treatment for 96 hrs. Viability was measured using a CellTiter-Glo assay. **k,** Nimbolide induces stronger cytotoxicity in UWB1 cells, compared to Olaparib. The cells were treated with the indicated compound for 96 hrs. Viability was measured using a CellTiter-Glo assay. **l, m,** Nimbolide overcomes intrinsic and acquired resistance to PARPi. **l,** HCC1937 cells were treated with nimbolide (1 μM) or PARPi (Olaparib and Rucaparib; 1 μM) for 96 hrs. Cell viability was measured using a CellTiter-Glo assay. **m,** UWB1 (SYr12) cells were treated with nimbolide or Olaparib for 96 hrs. Cell viability was measured using a CellTiter-Glo assay.

Our proteomic and biochemical studies indicate that PARylated-PARP1 is a novel substrate of RNF114 (Fig. 3). Based on these data, we hypothesized that nimbolide could represent a pharmacological approach to manipulating the RNF114-mediated ubiquitination and degradation of PARP1, and in doing so, to inducing PARP1 trapping. In this case, because nimbolide only occupies the E3 substrate recognition motif, but not the PAR-binding domain, of RNF114^40^, we hypothesized that the nimbolide-conjugated RNF114 could functionally mimic its RING domain mutations. Using the laser microirradiation assays, we found that nimbolide treatment resulted in profound PARP1 trapping (Fig. 4d). PARP1 trapping resulting from nimbolide treatment was very similar to that in the RNF114-*RING mutant cells. The kinetics of PARP1 trapping induced by nimbolide was also similar to that of Olaparib (a PARPi and a potent trapper of PARP1) (Fig. 4d).

Although both Olaparib and nimbolide trap PARP1, there is a unique distinction between these two classes of compounds. By blocking PARP1-mediated PARylation, Olaparib traps PARP1. In contrast, nimbolide inhibits RNF114 and thereby could result in the trapping of PARylated-PARP1. PAR polymers on PARylated-PARP1 are known to recruit many DNA repair factors (e.g., XRCC1), triggering the formation of a large protein complex involved in the repair of DNA SSBs^15^. We therefore hypothesize that nimbolide treatment could induce the trapping of not only PARylated-PARP1, but also other PAR-binding DNA repair proteins. To test this hypothesis, we performed laser microirradiation assays and found that, upon sensing DNA strand breaks, XRCC1 was rapidly recruited to the DNA lesions, through its PAR binding domains (Fig. 4e). Remarkably, XRCC1 accumulated and persisted at the DNA lesions in nimbolide-treated cells, likely owing to that PARylated-PARP1 was no longer removed in these cells. In contrast, Olaparib treatment blocked the formation of PAR chains, which completely abolished the recruitment of XRCC1 to the DNA lesions (Fig. 4e). Taken together, these data suggest that by targeting the E3 activity of RNF114, and subsequent proteasomal degradation of PARP1, nimbolide treatment causes potent PARP1 trapping. However, unlike conventional PARPi, nimbolide is a super trapper of not only PARP1, but also the PAR-dependent DNA repair factors.

We found that nimbolide-induced PARP1 trapping was abolished in RNF114-KO cells (Fig. 4c). Consistent with the role of trapped PARP1 in inducing cytotoxicity, we observed a dramatic increase of cell death in nimbolide-treated cells. However, these cytotoxic effects of nimbolide were largely abolished in RNF114-KO cells and PARP1-KO cells (Fig. 4g-i, Extended Data Fig. 3g). These data suggest that RNF114 is the primary mediator of the PARP1-trapping and cytotoxic activity of nimbolide.

Finally, we performed additional studies to further validate and probe the pharmacophore of nimbolide. Nimbolide contains a Michael acceptor moiety (Fig. 4a) which enables its covalent conjugation to RNF114 (e.g., via a Cys thiol group)^40^. We therefore synthesized several nimbolide analogs where we systemically manipulated this crucial enone moiety. The enone-reduced derivatives allylic alcohol **2** and ketone **3** were successfully obtained (Fig. 4b, Extended Data Fig. 4c and supplementary information). Consistent with the model where nimbolide covalently targets RNF114, these analogs (**2** and **3**) completely lost their cytotoxic activity (IC_50_ > 10 μM) against UWB1 cells (a *BRCA1*^mut^ ovarian cancer cell line, see more discussion below). The lactone opening analogs **4** and **5** (Extended Data Fig. 4c) were also found to be inactive. Therefore, we surmise that the enone and lactone moieties form the pharmacophore of nimbolide, which mediates its observed cytotoxic activity.

### Nimbolide treatment is synthetic lethal with *BRCA* mutations

It has been established that *BRCA1/2*-mutated cells are particularly sensitive to PARPi-induced trapping, and these cells are selectively killed by PARPi based on the “synthetic lethality” mechanism^6,13,14,41^. Indeed, we found that UWB1 cells were highly sensitive to nimbolide (IC_50_ = 0.3 μM). Furthermore, compared to the parental UWB1 cells, UWB1 cells reconstituted with *BRCA1* showed greatly reduced sensitivity to nimbolide (Fig. 4j). Consistent with the super trapping activity of nimbolide (which induces the trapping of not only PARP1, but also PAR-dependent DNA repair factors), nimbolide demonstrated superior cytotoxicity in UWB1 cells, compared to Olaparib (Fig. 4k). These data pointed to the synthetic lethality between nimbolide and *BRCA1* mutations. These results also suggest that *BRCA1* mutations (and potentially mutations of other genes in the homologous repair pathway) will serve as important predictive biomarkers for nimbolide sensitivity.

We also tested whether nimbolide acts synergistically with other DNA-damaging agents, including methyl methanesulfonate (MMS), Doxorubicin, and Temozolomide (TMZ). Compared to nimbolide alone, the combination of nimbolide with these agents showed significantly increased toxicity in UWB1 cells (Extended Data Fig. 4a). Furthermore, recent studies showed that “BRCAness” can also be induced by pharmacologically inhibiting key enzymes in the HR and DDR pathways^42^. Towards this, we treated UWB1 cells with nimbolide together with AZD6738 (an ATR inhibitor^43^), LY2603618 (a CHK1 inhibitor^44^), or SCH900776 (a CHK1 inhibitor^45^) (Extended Data Fig. 4b). Indeed, we found that nimbolide was also able to synergize with these agents to further enhance its cytotoxicity (Extended Data Fig. 4b).

### Nimbolide overcomes intrinsic and acquired resistance to PARP1 inhibitors

*BRCA1/2* mutations have been found in tumors originated in many different tissues, including breast, ovarian, prostate, and pancreas^46^. These mutations serve as excellent predictors for PARPi sensitivity. Although several PARPi have been approved for the treatment of breast and/or ovarian cancers with *BRCA* mutations, a significant fraction of the patients with *BRCA*^mut^ tumors showed *de novo* resistance, who failed to respond to these agents (intrinsic resistance)^47^. Because of the superior trapping activity of nimbolide (for both PARP1 and PAR-dependent DNA repair factors), we asked whether nimbolide is able to overcome intrinsic resistance to regular PARPi. HCC1937 is a *BRCA1*^mut^, triple-negative breast cancer cell line that is resistant to PARPi^48,49^. This cell line, however, was exquisitely sensitive to nimbolide even though it was resistant to regular PARPi (i.e., Olaparib and Rucaparib) (Fig. 4l).

Similar to other targeted therapies, those patients who showed initial response to PARPi often develop resistance, and relapsed disease is commonly observed. Thus, a strategy to overcome PARPi resistance is much needed to improve PARPi in order to achieve a more complete and durable response in the context of acquired resistance to PARPi. A previous study reported the UWB1 (SYr12) cells, which is a PARPi-resistant UWB1 clone derived from long-term culturing of the parental UWB1 cells in the presence of a PARPi (i.e., Olaparib). The resistance mechanism of these cells and the related clones have been ascribed to the transcriptionally rewired DNA damage response network^49^. We found that UWB1 (SYr12) cells showed sensitivity to nimbolide, but not Olaparib (Fig. 4m). Therefore, these data indicate that nimbolide is also able to kill tumors with acquired resistance to PARPi.

### Nimbolide triggers innate immune response and up-regulates PD-L1 expression

We recently showed that PARPi triggers innate immune signaling by PARP1 trapping-induced DNA damage response^21^. Because nimbolide induces potent PARP1 trapping and the subsequent DDR (Fig. 5a, b), we tested whether nimbolide could have any immunomodulatory roles. Immunofluorescence staining experiments showed that nimbolide treatment caused a significant accumulation of cytosolic double-stranded DNA (dsDNA) and micronuclei (Fig. 5c). Furthermore, we observed colocalization of Cyclic GMP-AMP synthase (cGAS) and cytosolic dsDNA in nimbolide-treated cells (Fig. 5c). cGAS is a critical sensor of cytosolic dsDNA. After the recognition of cytosolic dsDNA, cGAS generates the second messenger cGAMP (cyclic GMP-AMP), which then binds to and activate STING. This binding event results in the recruitment and activation of Tank-binding kinase I (TBK1). TBK1 phosphorylates a transcription factor Interferon Regulatory Factor 3 (IRF3), which leads to its nuclear translocation, and the activation of type I interferon (IFN) signaling^50–52^.

**Fig. 5.**
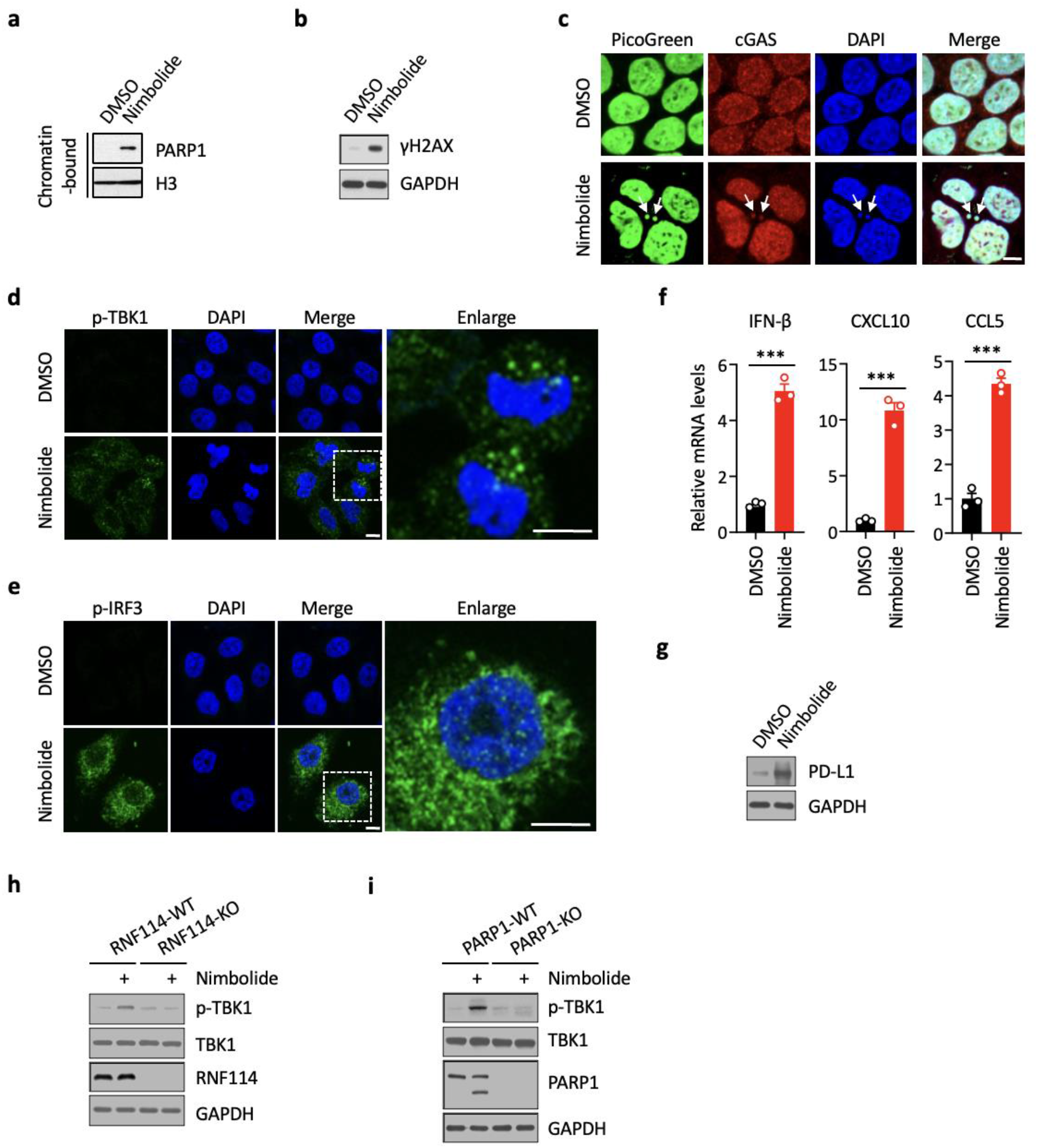
Nimbolide triggers an innate immune response and up-regulates PD-L1 expression. **a,** Nimbolide treatment induces PARP1 trapping. UWB1 cells were treated with or without nimbolide (1 μM for 48 hrs). The chromatin-bound fraction was isolated from these cells, and was subject to immunoblotting experiments using the indicated antibodies. **b,** Nimbolide treatment induces DNA damage response. HeLa cells were treated with or without nimbolide (1 μM for 48 hrs). The cell lysates were subject to immunoblotting experiments using the indicated antibodies. **c,** Nimbolide treatment induces the formation of cytosolic dsDNA and micronuclei. HeLa cells were treated with or without nimbolide (1 μM for 48 hrs). DAPI (blue) was used to visualize the nuclei. Arrows indicate cytosolic dsDNA and micronuclei. Scale Bars, 10 μm. **d,** Nimbolide treatment induces the phosphorylation of TBK1. HeLa cells were treated with or without nimbolide (1 μM for 48 hrs). The level of pS172 TBK1 (p-TBK1, Green) was detected using the immunofluorescence assay. DAPI (blue) was used to visualize the nuclei. Scale Bars, 10 μm. **e,** Nimbolide treatment induces the phosphorylation of IRF3. HeLa cells were treated with or without nimbolide (1 μM for 48 hrs). The level of pS396 IRF3 (p-IRF3, Green) was detected using the immunofluorescence assay. DAPI (blue) was used to visualize the nuclei. Scale Bars, 10 μm. **f,** qRT-PCR analyses of IFN-β, CXCL10, or CCL5 in HeLa cells treated with or without nimbolide (1 μM for 48 hrs). **g,** Nimbolide treatment induces the expression of PD-L1. UWB1 cells were treated with or without nimbolide (1 μM for 48 hrs). PD-L1 expression was detected using the immunoblot assay. **h,** RNF114-KO abrogates the nimbolide-induced TBK1 phosphorylation. Control (RNF114-WT) and RNF114-KO HeLa cells were treated with or without nimbolide (1 μM for 48 hrs). The whole cell lysates were subject to immunoblot experiments using the indicated antibodies. **i,** PARP1-KO abrogates the nimbolide-induced TBK1 phosphorylation. Control (PARP1-WT) and PARP1-KO HeLa cells were treated with or without nimbolide (1 μM for 48 hrs). The whole cell lysates were subject to immunoblot experiments using the indicated antibodies.

We found that phosphorylation of TBK1 (p-TBK1), a key downstream effecter of cGAS-STING pathway, was dramatically upregulated in nimbolide-treated cells (Fig. 5d). Furthermore, nimbolide treatment induced the activation and nuclear translocation of p-IRF3, suggesting the activation of cGAS-STING pathway in these cells (Fig. 5e). To further assess the activation of the cGAS-STING pathway, we examined the mRNA levels of a number of downstream target genes in cGAS-STING pathway. Indeed, the mRNA levels of IFN-β, CXCL10 and CCL5 were dramatically increased in nimbolide-treated cells (Fig. 5f). Finally, consistent with the superior trapping activity of nimbolide, it was able to induce stronger activation of the cGAS-STING pathway (as shown by higher levels of p-TBK1), compared to other PARPi (i.e., Olaparib) (Extended Data Fig. 5a). Collectively, these results demonstrate that nimbolide induces the accumulation of cytosolic dsDNA, which then activates the cGAS-STING-TBK1-IRF3 innate immune signaling.

A critical downstream target of the cGAS-STING pathway is programmed death-ligand 1 (PD-L1), a major ligand of PD-1^53^. The binding of PD-L1 to the immune checkpoint molecule PD-1 transmits an inhibitory signal to reduce the proliferation of antigen-specific T cells^54^. Recent studies suggested that PD-L1 expression is regulated by PARPi^55^. Since nimbolide treatment induces PARP1 trapping, DNA damage, and innate immune response, we tested whether PD-L1 is also regulated by nimbolide treatment. Indeed, we found that the expression of PD-L1 was greatly elevated in nimbolide-treated UWB1 cells (Fig. 5g). Consistent with the superior trapping activity of nimbolide, it was able to induce stronger expression of PD-L1 compared to Olaparib (Extended Data Fig. 5b). Nimbolide failed to induce TBK1 activation in RNF114-KO or PARP1-KO cells, indicating the specificity of nimbolide in the context of its immunomodulatory roles (Fig. 5h, i). Together, these data suggest that nimbolide activates the innate immune response and up-regulates PD-L1 expression. These results raise the hypothesis that by inducing the activation of the cGAS-STING pathway, nimbolide could synergize with immune checkpoint inhibitors.

## Discussion

Despite the tremendous progresses of PARPi in the clinic, the mechanism of action of PARPi remains incompletely understood. All FDA-approved PARPi possess two distinct, yet interconnected activities, namely PARP1 inhibition and PARP1 trapping^17^. It was initially thought that PARPi kills tumor cells simply by inhibiting the catalytic activity of PARP1, and thus blocking PARylation-dependent DDR signaling^15^. More recent studies showed that, upon the treatment of PARPi, PARP1 is retained at the DNA damage site for an extended time (termed “trapping”), resulting in the formation of a trapped DNA-PARP1-PARPi complex^6,18^. The trapped PARP1 causes the collapse of the replication fork, and is known to be highly toxic to cells^6^. Although various studies have provided compelling evidence pointing to PARP1 trapping as likely a main contributor to the anti-tumor activity of PARPi (e.g., the induction of DDR, cytotoxicity and innate immune signaling)^18,21–24^, the molecular nature of PARP1 trapping remains poorly understood.

To identify the regulatory factors involved in the PARylation dependent DNA damage response, we herein performed a chromatin localization screen. In this screen, we used quantitative proteomic experiments to identify proteins that become enriched or depleted in the chromatin fraction, upon the treatment of genotoxic agents. From these experiments, we garnered unanticipated insights into the molecular underpinnings of PARP1 trapping. Specifically, in response to genotoxic stress, we found that a poorly studied ubiquitin E3 ligase, RNF114, becomes enriched in the chromatin fraction. Importantly, the recruitment of RNF114 to DNA lesions is mediated by the interaction between PAR chains and the C-terminal PBZ domain of RNF114. Besides the role as a scaffold to recruit RNF114, PAR chains also stimulate the E3 activity of RNF114. Using an IP-MS approach, we subsequently identified PARylated-PARP1 as a novel RNF114 substrate, and RNF114 specifically targeted PARylated-PARP1 for ubiquitin-proteasomal degradation.

From these functional studies, we identified a novel cross-talk mechanism between PARylation and ubiquitination that is mediated by the RNF114. Importantly, the coupling between PARylation and RNF114-mediated ubiquitination and degradation of PARP1 has profound implications in the context of PARP1 trapping. Upon binding to DNA lesions, PARP1 is rapidly activated to promote its own PARylation^56,57^. It has been proposed that PARP1 could be dissociated from DNA, owing to the steric hindrance and charge repulsion introduced by the PAR polymers^58^. PARPi treatment prevents PARP1 auto-PARylation and causes it to be trapped at DNA lesions.

Our results suggest that, in response to genotoxic stimuli, RNF114 is recruited to the DNA damage sites by its binding to PARylated-PARP1. RNF114 then degrades PARylated-PARP1, which provides a novel ubiquitination-dependent mechanism to remove PARylated-PARP1 from DNA lesions. PARPi inhibit the PARylation of PARP1, and thus prevent the recruitment of RNF114 to the DNA lesions. This subsequently blocks the ubiquitination and degradation of PARP1, leading to PARP1 trapping. Consistent with this model, we observed potent PARP1 trapping in cells expressing the RNF114-*RING mutant. The binding of the RNF114-*RING mutant could shield PARylated-PARP1 and therefore prevent its removal by other factors.

In the context of PARP1 removal from the chromatin, RNF114-mediated degradation is likely complementary with respect to the PAR-repulsion model. The in vivo metabolism of PARylation is highly dynamic, which is under the tight regulation of not only PARP1, but also many PAR-catabolizing enzymes (e.g., Poly-ADP-ribose glycohydrolase, PARG)^59^. It is likely that, under these conditions, the steric hinderance-introduced by PARylation (e.g., high local concentrations of PAR polymers) is still required for the release of PARP1 from DNA^60–62^. It is therefore possible that the RNF114-mediated degradation pathway functions coordinately with these mechanisms to drive the removal of PARP1 from the DNA damage site.

Nimbolide is a limonoid natural product that is derived from the Neem tree^35^. Although this compound has previously been shown to possess anti-cancer activity, its exact mechanism of action was poorly characterized^63^. Using a chemoproteomic approach, a recent study showed that nimbolide covalently modifies RNF114, and in doing so, prevents its E3 substrate engagement^40^. We identified, in the current study, PARylated-PARP1 as a novel substrate of RNF114, and blockage of this pathway prevents the ubiquitination and degradation of PARP1 (Fig. 6). Based on these findings, we hypothesized that by targeting RNF114, nimbolide treatment could serve as a pharmacological approach to manipulate the degradation of PARP1. Upon nimbolide treatment, the ability of RNF114 to recognize PARylated-PARP1 could be compromised, which could lead to impaired ubiquitination and proteasomal degradation of PARP1. Mechanistically, nimbolide mimics the RNF114-*RING mutant to introduce the dominant negative effect, which results in potent PARP1 trapping.

**Fig. 6.**
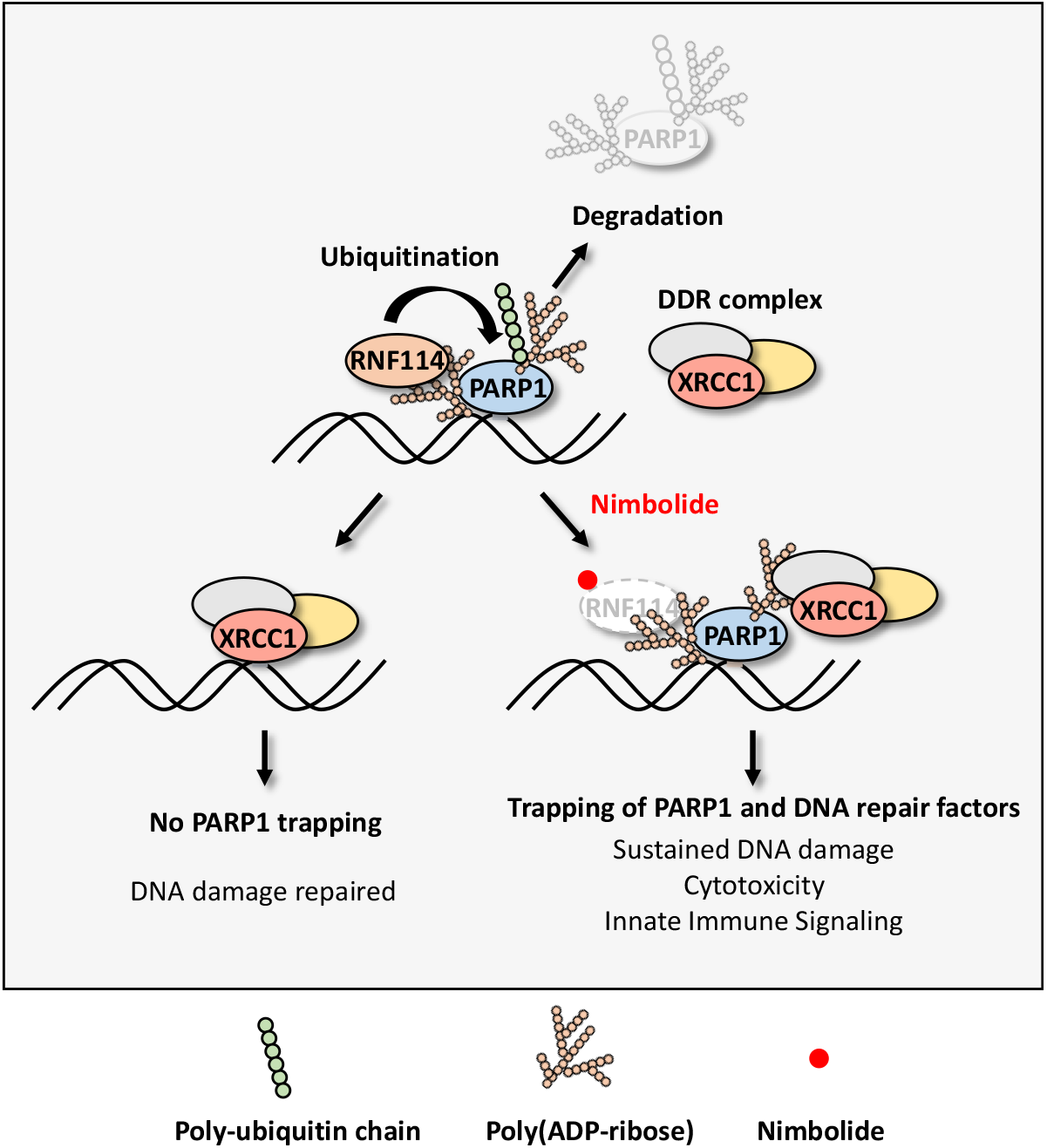
A schematic model of the nimbolide-induced trapping of PARylated-PARP1 and PAR-dependent DNA repair factors.

Although both PARPi and nimbolide induce PARP1 trapping, a key difference between these two classes of compounds is that PARPi traps PARP1, whereas nimbolide traps PARylated-PARP1 (Fig. 6). In this case, PARPi cause PARP1 trapping by blocking its auto-PARylation, and also by allosterically enhancing its DNA binding^61^. In contrast, nimbolide treatment prevents the removal of PARylated-PARP1 from DNA lesions, leading to potent trapping of PARylated-PARP1 (Fig. 6). Protein-linked PAR polymers are known to recruit many proteins that bear PAR-binding domains. As an example, in response to genotoxic stress, PAR polymers function as a scaffold to recruit a protein called XRCC1, triggering the formation of a large protein complex involved in the repair of DNA SSBs^15^. Indeed, we found that PARPi treatment completely abolished the recruitment of XRCC1 to DNA lesions, whereas potent retention of XRCC1 was observed in nimbolide-treated cells. These data again suggest that RNF114 inhibition impairs the removal of PARylated-PARP1 from the DNA damage site, leading to the subsequent retention of both PARylated-PARP1 and PAR-dependent DNA repair factors.

Although the various FDA-approved PARPi inhibit the catalytic activity of PARP1 with similar potency, they differ in their allosteric binding to PARP1, leading to the unequal PARP1-trapping activities^61^. Importantly, it has been shown that DDR, cytotoxicity and innate immune responses in PARPi-treated cells are all positively correlated with the degree of PARP1 trapping induced by the specific PARPi^21,23^. Because nimbolide traps not only PARylated-PARP1, but also the PAR-dependent DNA repair factors, we then examined the anti-cancer activity of nimbolide (Fig. 6). First, we found that, similar to other PARPi, nimbolide treatment is synthetically lethal with respect to *BRCA1* mutations. Specifically, we showed that nimbolide demonstrated enhanced cytotoxicity against UWB1 cells, compared to UWB1 cells reconstituted with *BRCA1* (Fig. 4j). These results suggest that mutations of *BRCA1* (and likely *BRCA2* and other genes in the homologous recombination pathway) will serve as important predictive biomarkers for nimbolide sensitivity. Second, we showed that the unique trapping activity of nimbolide can be translated into its distinct advantages over PARPi in overcoming PARPi resistance (both intrinsic and acquired resistance). Specifically, we showed that both HCC1937 (a *BRCA1*^mut^ breast cancer cell line that is intrinsically resistant to regular PARPi) and UWB1 (SYr12) (a *BRCA1*^mut^ ovarian cancer cell line that has developed acquired resistance to PARPi) remain exquisitely sensitive to nimbolide (Fig. 4l, m). Third, nimbolide was able to act synergistically with DNA-damaging agents and DNA repair machinery inhibitors (e.g., ATR and CHK1 inhibitors) (Extended Data Fig. 4a, b). We showed that by inducing PARP1 trapping and DDR, nimbolide triggers the activation of innate immune signaling. This leads to the elevated expression of immunomodulatory proteins, including PD-L1.

Finally, PARG is known to be a highly active enzyme that removes the PAR chains from PARP1 and other PARP1 substrates^59^. This suggests that by enhancing the PARylation level, PARG inhibitors could further promote the trapping of PAR-dependent DNA repair factors, under RNF114-suppressed conditions (e.g., nimbolide-treated cells). We therefore examined the potential synergistic effects between PARG inhibitors and nimbolide. Indeed, compared to nimbolide treatment alone, the combination of nimbolide with PDD00017273 (a highly potent PARG inhibitor^64–66^) showed increased toxicity in UWB1 cells (Extended Data Fig. 6).

Collectively, we showed that enhanced trapping of PARP1 and PARylated DNA repair factors induced by nimbolide treatment can be translated into distinct advantages compared to NAD^+^-competitive PARPi. The full therapeutic potential of nimbolide and its analogs as a monotherapy, and its combination with other DNA-damaging agents and immune checkpoint inhibitors warrants future studies.

In summary, our studies suggest that in response to genotoxic stress, PARP1 is recruited to DNA lesions, and becomes auto-PARylated (Fig. 6). RNF114 is then recruited, in a PAR-dependent manner, to the DNA damage site, to catalyze the ubiquitination and subsequent proteasomal degradation of PARP1. By targeting the substrate recognition domain of RNF114, nimbolide treatment blocks PARP1 degradation, leading to PARP1 trapping. Importantly, unlike regular PARPi, nimbolide treatment induces the trapping of not only PARylated-PARP1, but also PAR-dependent DNA repair factors. We then showed that nimbolide is synthetic lethal with respect to *BRCA* mutations, and it was able to kill cancer cells with intrinsic and acquired resistance to PARPi. Finally, we demonstrated that nimbolide treatment is synergistic with other DNA damaging agents, activates innate immune response and up-regulates PD-L1 expression. Our results point to the exciting possibility of therapeutically targeting the RNF114-dependent PARP1 degradation pathway for the treatment of *BRCA*^mut^ cancers.

## Acknowledgments

We thank Dr. Lee Zou for providing the UWB1, UWB1+BRCA1 and UWB1 (SYr12) cells; Dr. Xiaochun Yu for providing the PARP1-GFP and XRCC1-GFP plasmids. This work was supported in part by grants from the Welch Foundation (I-2010-20190330 to T.Q. and I-1800 to Y.Y.), and UT Southwestern Eugene McDermott Scholarship (T.Q.) and NIH (5R35GM134883, 1R01NS122533 and 1R21CA261018 to Y.Y., R01GM141088 to T.Q.).

## Author contributions

P.L., Y.Z., C.K., X.W. and Y.Y. performed the biology-related experiments; H.D., H.D. and T.Q. synthesized the nimbolide analogs. Y.Y. and T.Q. designed and supervised the project; all authors wrote the paper;

## Competing interests

Provisional patents have been filed on some aspects of the work described in this manuscript. Y.Y. and T.Q. are co-founders and shareholders of ProteoValent Therapeutics;

## Data and materials availability

Experimental procedures, alternative synthetic route, optimization data, ^1^H NMR spectra, ^13^C NMR spectra and MS data are available in the Supplementary Materials.

## Materials and Methods

### Materials

The antibodies used in this study were obtained from commercial sources, including anti-PAR (Trevigen; 4335-MC-100), anti-GAPDH (Thermo Fisher Scientific; MA1-16757), anti-Flag-tag (Thermo Fisher Scientific; MA1-91878), anti-GST (CST; 2622S), anti-RNF114 (Santa Cruz; sc-514747), anti-Histone (CST; 4499S), anti-Ubiquitin (Santa Cruz; sc-8017), anti-PARP1 (CST; 9542S), anti-γH2AX (CST; 9718S), anti-p-TBK1 (CST; 5483S), anti-TBK1 (CST; 38066S), anti-p-IRF3 (CST; 29047S), anti-IRF3 (CST; 11904S), anti-cGAS (CST; 15102S), anti-STING (CST; 13647S) and anti-PD-L1 (CST; 13684S). nimbolide was purchased from Sigma-Aldrich (SMB00586). PARP inhibitors: Rucaparib (Selleck; S4948), Olaparib (Selleck; S1060), Talazoparib (Selleck; S7048) were purchased from the indicated sources. All other reagents including MG132 (Selleck; S2619), H_2_O_2_ (Thermo Fisher Scientific; H325), MMS (Sigma-Aldrich; 129925), Doxorubicin (Sigma-Aldrich; D1515), Temozolomide (Sigma-Aldrich; T2577), AZD6738 (Cayman; 21035), LY2603618 (Apexbio; A8638), SCH900776 (Cayman; 18131) were obtained from commercial sources.

### Cell culture

All the cells were purchased from ATCC and cultured according to the directions from ATCC. HCT116 and HeLa cells were maintained in the high-glucose DMEM medium supplemented with 10% FBS. HCC1937 cells were maintained in RPMI1640 (ATCC) supplemented with 15% FBS. UWB1, UWB1+BRCA1, and UWB1 (SYr12) cells were gifts from Dr. Lee Zou (Harvard Medical School). UWB1 and UWB1+BRCA1 cells were maintained in RPMI1640 (ATCC) and MEGM bullet kit (1:1; Lonza) with 3% FBS. UWB1 (SYr12) cells were maintained in RPMI1640 and MEGM bullet kit with 3% FBS and 1 μM PARPi (Olaparib; Selleck)^49^.

### Plasmid and construction

The RNF114 clone was obtained from the Center for Human Growth and Development of UTSW. The RNF114 cDNA was subcloned into the pcDNA3 (addgene), pCDNA5-ZZ-TEV-Flag vector (addgene), or pGEX-4T-3 (addgene) vectors for transient transfections. Besides, the RNF114 cDNA was transferred into the plenti-6.3-V5-Dest vector (Thermo Fisher) for the subsequent construction of stable cell lines. Various site mutations of RNF114 were introduced using standard site-directed mutagenesis techniques. RNF114 CRISPR KO constructs were made according to a previously described protocol *(Nature Methods* volume 11, pages783–784 (2014)). Basically, three top ranked sgRNAs were chosen (RNF114 sg1F: CACCGGTGTACGAGAAGCCGGTAC; RNF114 sg1R: AAACGTACCGGCTTCTCGTACACC; RNF114 sg2F: CACCGGACACGTGAAGCGTCCTAG; RNF114 sg2R: AAACCTAGGACGCTTCACGTGTCC; RNF114 sg3F: CACCGCACGTGTCCCGTGTGCTTAG; RNF114 sg3R: AAACCTAAGCACACGGGACACGTGC), and they were subcloned into *LentiCRISPR* v2 vector (Addgene). All the mutant constructs were confirmed by DNA sequencing analysis. PARP1-GFP and XRCC1-GFP plasmids were gifts from Dr. Xiaochun Yu (City of Hope).

### Construction of stable cell lines

The plenti or pLKO.1 construct (8 μg), VSVG (6 μg), and delta8.9 (6μg) were co-transfected into HEK293TD cells in 10cm dishes with Lipo-2000 (Sigma). The medium was changed 6 h after transfection. Viruses were collected twice at 24 h and 48 h after transfection respectively, and then combined together. Subsequently, 3 mL of virus was added to each well of HCT116 or HeLa cells in 6-well plates with Polybrene (8 μg/ml). After splitting the cells once, HCT116 or HeLa cells were infected with the previously collected virus again using the same procedure. The culture medium was replaced after 48 h with a fresh growth medium containing 2 μg/ml blasticidin or puromycin. To generate the RNF114 CRISPR KO cell lines, we performed single cell clone selection after recombinant lentiviruses infection (the viruses were produced by transfecting all three *LentiCRISPR* v2-sgRNF114 plasmids together). The clones that were completely depleted of the RNF114 protein were chosen for future experiments.

### Sample preparation and the mass spectrometry-based chromatin re-localization screen

To analyze how chromatin proteins respond to genotoxic stress, HCT116 cells were pre-treated with Talazoparib (1 μM for 1 h), which was followed by the treatment with H_2_O_2_ (2 mM for 5 min) or MMS (0.01% for 1 h) as indicated. Cells were washed with cold 1×PBS and the chromatin-bound proteins were extracted using the subcellular fractionation kit (Thermo Fisher). Protein concentrations were measured by the BCA assay kit (Thermo Fisher). For the TMT experiments, proteins were reduced with dithiothreitol (2 mM for 10 min) and alkylated with iodoacetamide (50 mM for 30 min) in the dark. Proteins were then extracted by methanol/chloroform precipitation and were washed by ice-cold methanol. Protein pellets were re-dissolved in 400 μl of an 8 M freshly prepared urea buffer (50 mM Tris-HCl and 10 mM EDTA, pH 7.5). The proteins were digested by Lys-C at a 1:100 enzyme/protein ratio for 2 h at room temperature, followed by trypsin digestion at a 1:100 enzyme/protein ratio overnight at room temperature. Peptides were desalted with Oasis HLB cartridges and were resuspended in 200 mM HEPES, pH 8.5. For each sample, 100 μg of peptides were reacted with the corresponding amine-based TMT six-plex reagents (Thermo Fisher) for 1 h at RT. The labeling scheme was as following: 126: control, 127: H_2_O_2_, 128: MMS, 129: Talazoparib, 130: Talazoparib and H_2_O_2_ and 131: Talazoparib and MMS. These actions were quenched with a hydroxylamine solution and the peptide samples were combined.

The TMT samples were desalted and were fractionated by basic pH reversed phase HPLC on a ZORBAX 300 Extend-C18 column. Buffer A was 10 mM ammonium formate in water, pH 10.0. A gradient was developed at a flow rate of 0.2 ml/min from 0% to 70% buffer B (1 mM ammonium formate, pH 10.0, 90% acetonitrile). Seventeen fractions were collected, which were lyophilized, desalted and analyzed by LC-MS/MS experiments as described previously^23,27,67^.

### Laser microirradiation assays

Cells grown on 35-mm glass-bottomed culture dishes (Mattek) were transfected with the GFP-tagged RNF114, GFP-PARP1, or GFP-XRCC1 plasmid for 24 h. After the compound treatment as indicated in each experiment, laser microirradiation was performed using a Zeiss LSM 780 inverted confocal microscope coupled with the MicroPoint laser illumination and ablation system (Photonic Instruments). The GFP fluorescence at the laser line was recorded at the indicated time points and was then analyzed with the ImageJ software.

### Immunoblot analysis

Cells were collected and washed once with cold 1×PBS. Then, cells were lysed with the 1% SDS lysis buffer (1% SDS, 10 mM HEPES, pH 7.0, 2 mM MgCl_2_ and 500 U universal nuclease). Protein concentrations were measured by the BCA assay kit (Thermo Fisher). The same amount of protein was loaded onto an SDS-PAGE gel. After electrophoretic separation, proteins were transferred to a NC (nitrocellulose) membrane (GE Healthcare). For the dot-immunoblot analysis, samples were loaded directly to the NC membrane. The membrane was blocked and was then blotted with the primary antibodies overnight at 4 °C, which was followed by incubation with the secondary antibody for 1 h at room temperature. The blots were developed using enhanced chemiluminescence and were exposed on autoradiograph films.

### Co-immunoprecipitation analysis

Cells were collected and lysed in the IP lysis buffer (50 mM Tris, pH 7.4, 1 mM EDTA, 150 mM NaCl, 0.5% NP-40, and 1×protease inhibitor cocktail). After incubation for 1 h at 4 °C, cell lysates were centrifuged at 14,000 g for 10 min at 4 °C. The supernatants were transferred and incubated with the corresponding agarose beads overnight at 4 °C. The beads were washed three times with the IP wash buffer, and the immunocomplexes were eluted from the beads by boiling at 95 °C for 30 min and were subject to immunoblot analysis. The TAP-IP-MS experiments were performed according to a previously described protocol^68^.

### Immunofluorescence microscopy

After the treatment with DMSO or nimbolide, HeLa cells were washed once with 1×PBS, and then, fixed with 4% paraformaldehyde for 20 min at room temperature followed by three times wash with 1×PBS. The cells were permeabilized with 0.25% Triton X-100 in 1×PBS for 10 min and blocked with 1×PBS containing 2% BSA for1 h. Fixed cells were incubated with primary antibodies at 4 °C overnight, followed by the incubation with the fluorescent secondary antibody for 1 h at room temperature (RT). Cells were washed three times with 1×PBS for 5 min and stained with DAPI (ThermoFisher Scientific) for 2 min. Cells were washed with 1×PBS and mounted with the FluorSave reagent (Millipore). The fluorescence images were then collected with a Zeiss LSM 880 Airyscan inverted confocal microscope.

### qRT-PCR

The cells with the treatment of DMSO or nimbolide were lysed with TRIzol (Thermo Fisher). Then, the total RNA was extracted according to the manufacturer’s protocol and subject to reverse transcription with SuperScript® III One-Step RT-PCR System (Thermo Fisher). The qRT-PCR experiments were performed with a Power SYBR® Green PCR Master Mix (Thermo Fisher) and specific primers listed as following: GAPDH, sense: ACAACTTTGGCATTGTGGAA, anti-sense:GATGCAGGGATGATGTTCTG; IFN-β, sense: AGCTGAAGCAGTTCCAGAAG, anti-sense: AGTCTCATTCCAGCCAGTGC; CXCL10, sense: GGCCATCAAGAATTTACTGAAAGCA; anti-sense: TCTGTGTGGTCCATCCTTGGAA; CCL5, sense: ATCCTCATTGCTACTGCCCTC; anti-sense: GCCACTGGTGTAGAAATACTCC.

### *In vitro* PARylation assays

PARP1 (500 ng, Tulip Biolabs), sheared Salmon Sperm DNA (100 ng, Thermo Fisher) and NAD^+^ (500 μM) were incubated in the reaction buffer (50 mM Tris, pH 7.5, 4 mM MgCl_2_, 20 mM NaCl and 250 μM DTT) at RT for 1 h. Reactions were terminated by the SDS loading buffer and the samples were subject to immunoblot analyses by an anti-PAR antibody.

### *In vitro* ubiquitination assays

To measure the auto-ubiquitination of RNF114, UBE1 (50 nM, Boston Biochem), UBE2D1 (50 nM, Boston Biochem) and Ubiquitin (200 mM, Boston Biochem) were incubated with the recombinant RNF114 (1 μg) at 37°C in the ubiquitination reaction buffer (50 mM Tris-Cl, pH 7.5, 2.5 mM MgCl_2_, 2 mM DTT, 2 mM ATP). Reactions were terminated by SDS loading buffer and boiled. The supernatant samples were subject to immunoblot analysis by an anti-ubiquitin antibody. For the *in vitro* PARylated-PARP1 ubiquitination assays, PARP1 or PARylated-PARP1 (350 ng) was incubated together with UBE1 (50 nM, Boston Biochem), UBE2D1 (50 nM, Boston Biochem), Ubiquitin (200 μM, Boston Biochem) and recombinant RNF114 (1μg) at 37°C in the ubiquitination reaction buffer as above. Reactions were terminated by the SDS loading buffer and boiled. The supernatant samples were subject to immunoblot analyses by the anti-ubiquitin and anti-PAR antibodies.

### Colony formation assays

HCT116 control (RNF114-WT) or RNF114-KO cells were seeded into 6-cm dishes (about 1000 cells per dish). The cells were treated with or without H_2_O_2_ (2 mM for 5 min) and followed by 14 days culture. The viable cells were fixed by methanol and were stained with crystal violet.

### Cell viability measurement

Cells were plated into 96-well plates at densities of 1000 cells/well. Next day, cells were treated as indicated. Cell viability was measured using the CellTiter-Glo assay (Promega) according to the manufacturer’s instructions. Briefly, after incubation, room temperature CellTiter-Glo reagent was added 1:1 to each well, and the plates were incubated at room temperature for 2 min. Luminescence was measured with the Synergy HT Multi-Detection Microplate Reader and was normalized against control cells.

### Statistics

All of the statistical analyses (*t*-tests) were performed using the GraphPad Prism software (v.8). Data were derived from the average of three biological replicate experiments and were presented as the mean ±SEM. ^*^*P* < 0.05, ^**^*P* < 0.01 and ^***^*P* < 0.001.

## Supplementary figures

**Extended Data Fig. 1.**
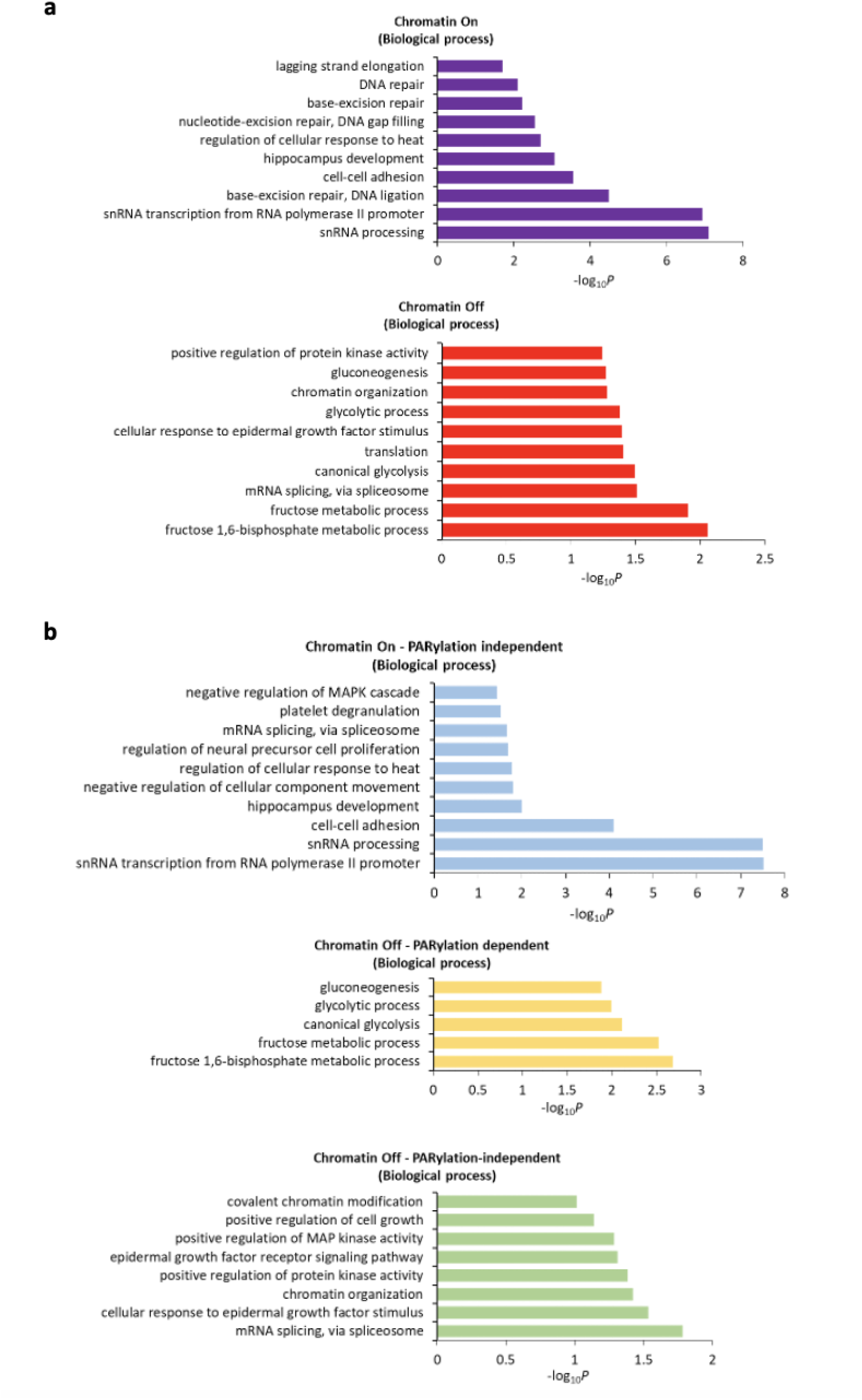
GO analysis of the Chromatin-On and -Off proteins. **a,** GO analysis of the Chromatin-On and -Off proteins. These proteins include all the proteins (i.e., PARylation-dependent and PARylation-independent). **b,** GO analysis of the Chromatin-On-PARylation independent, Chromatin-Off-PARylation dependent, and Chromatin-Off-PARylation independent proteins.

**Extended Data Fig. 2.**
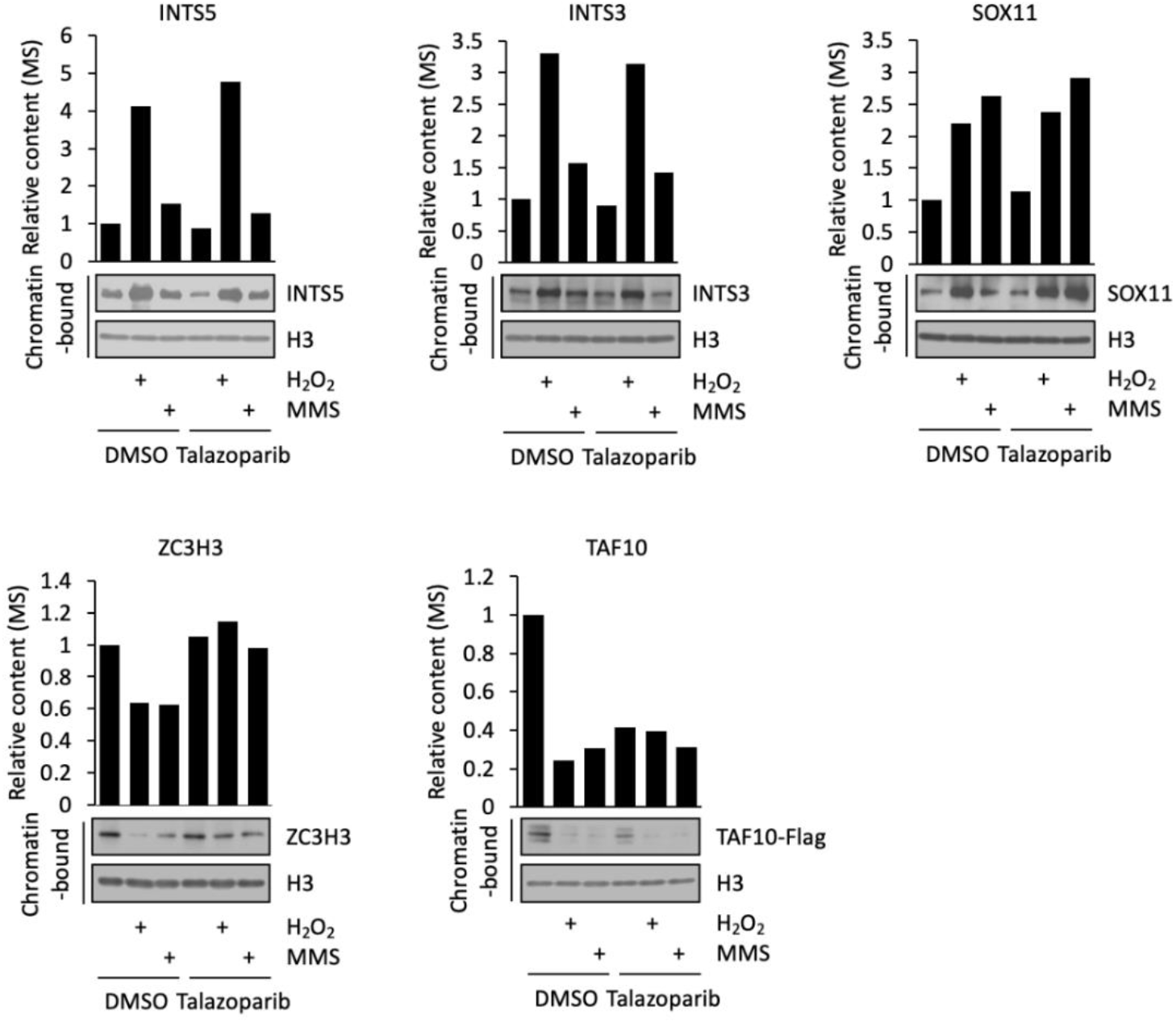
Validation of DDR-induced re-localization events. The abundance of INTS5, INTS3, SOX11, ZC3H3, and TAF10 in the chromatin fraction as measured in the proteomic (upper panel) and immunoblot experiments (lower panel).

**Extended Data Fig. 3.**
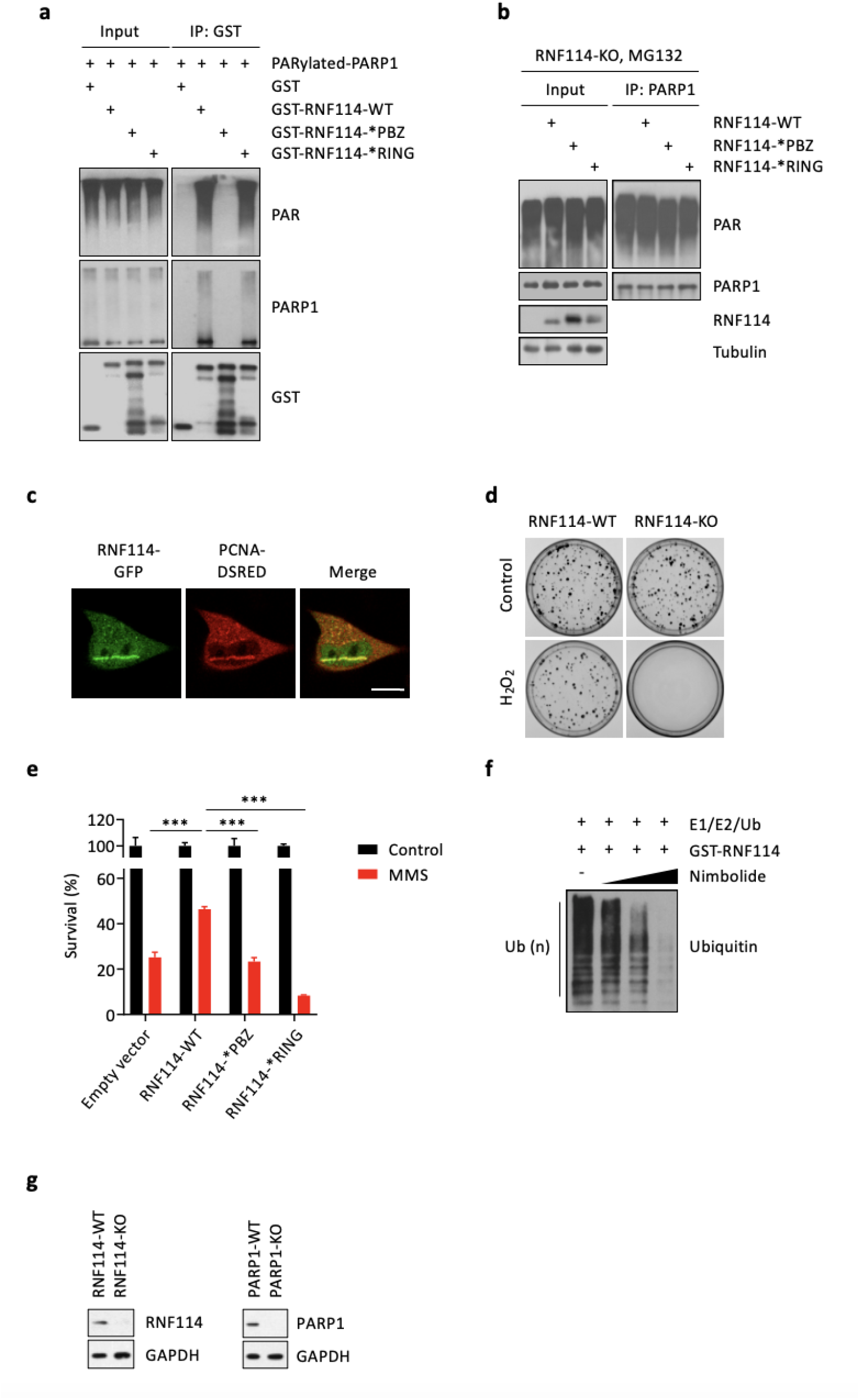
RNF114 targets PARylated-PARP1 for ubiquitin-proteasomal degradation. **a,** RNF114 interacts with PAR. Immunoblot analyses of the interaction between PARylated-PARP1 and RNF114 *in vitro*. The recombinant GST-RNF114-WT, GST-RNF114-*PBZ mutant, or GST-RNF114-*RING mutant was incubated with PARylated-PARP1. The samples were subject to glutathione-based enrichment (for the isolation of GST and GST-fusion proteins). **b,** PARylated-PARP1 is not degraded by RNF114 in proteasome inhibition. RNF114-KO cells were reconstituted with RNF114-WT, RNF114-*PBZ mutant, or RNF114-*RING mutant. These cells were pre-treated with MG132 (10 μM for 6 h) and were then treated with H_2_O_2_ (2 mM for 5 min). PARP1 was isolated using immunoprecipitation and was subject to immunoblot analyses using the indicated antibodies. **c,** RNF114 is co-localized with PCNA during DNA damage response. Staining of RNF114-GFP (Green) and PCNA-DSRED (Red) during laser microirradiation-induced DNA damage. Scale Bars, 10 μm. **d,** RNF114 is involved in DNA damage response. Control (RNF114-WT) and RNF114-KO HCT116 cells were treated with or without H_2_O_2_ (2 mM for 5 min). Cell viability was measured using the colony formation assay. **e,** Mutation of the RING domain in RNF114 renders cells susceptible to genotoxic stress. RNF114-KO HCT116 cells were reconstituted with RNF114-WT, RNF114-*PBZ mutant, or RNF114-*RING mutant. The cells were treated with MMS (1 mM for 9 h). Cell viability was measured using the CellTiter-Glo assay. **f,** Nimbolide blocks the auto-ubiquitination of RNF114 in an in vitro ubiquitination assay. Where indicated, GST-RNF114 was incubated with E1/E2/ubiquitin in the presence of increasing concentrations of nimbolide (0.1 μM, 0.5 μM or 1 μM). **g,** Generation of the RNF114-KO and PARP1-KO HeLa cells. RNF114 or PARP1 was deleted in HeLa cells using the CRISPR-Cas9 system. Whole cell lysates were subject to immunoblot analyses using the indicated antibodies.

**Extended Data Fig. 4.**
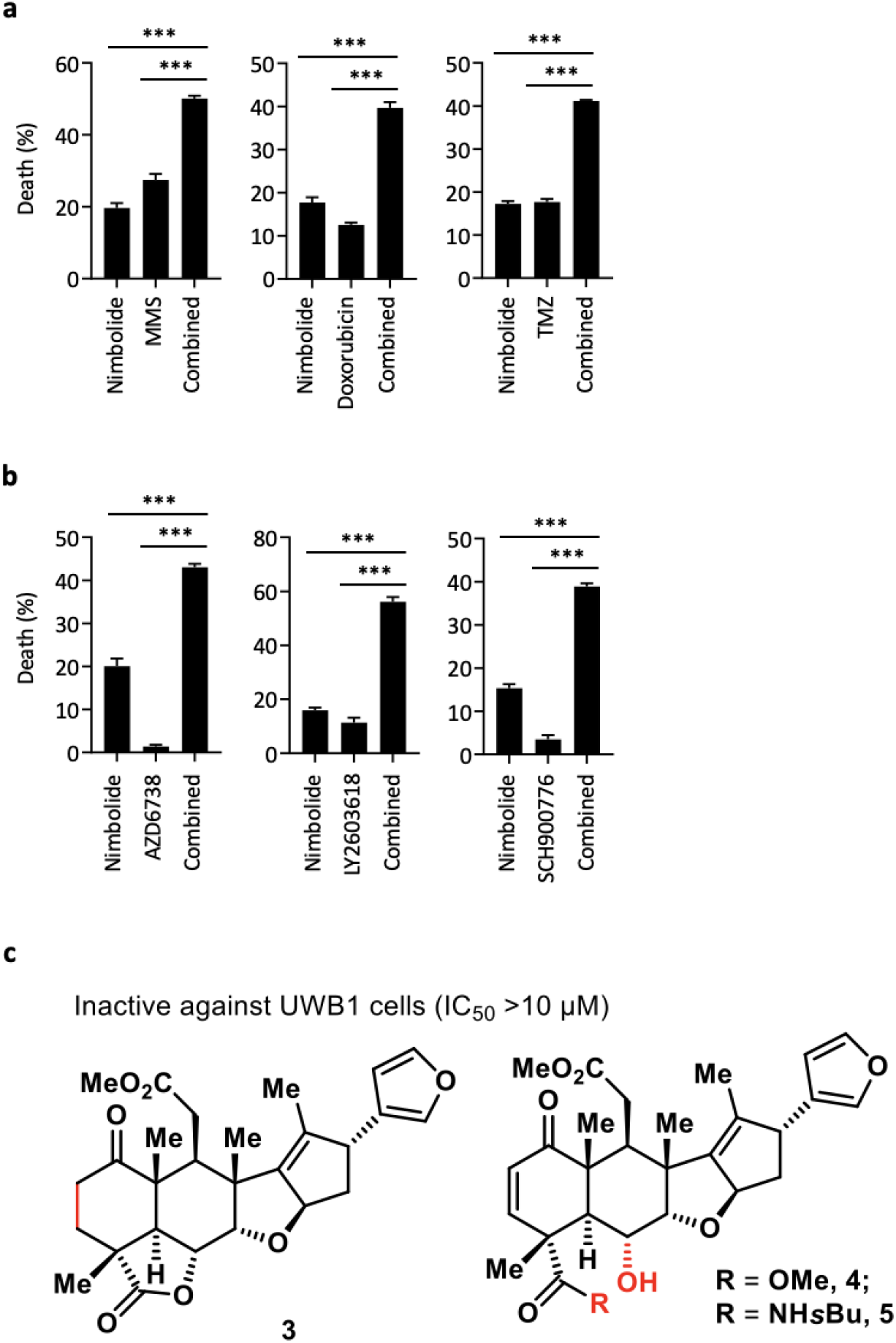
Nimbolide synergizes with DNA damaging agents. **a,** The synergistic effect between nimbolide and DNA damaging agents. UWB1 cells were treated with either nimbolide (0.25 μM) alone, or nimbolide in combination with various DNA-damaging agents (MMS (10 μM), Doxorubicin (0.01 μM), and Temozolomide (TMZ, 10 μM)) for 96 hrs. **b,** The synergistic effects between nimbolide and inhibitors of the DNA damage repair machinery. UWB1 cells treated with either nimbolide (0.25 μM) alone, or nimbolide in combination with (AZD6738 (0.1 μM), LY2603618 (0.1 μM), or SCH900776 (0.1 μM)) for 96 hrs. **c,** Structure of the various nimbolide analogs. These compounds were inactive against UWB1 cells.

**Extended Data Fig. 5.**
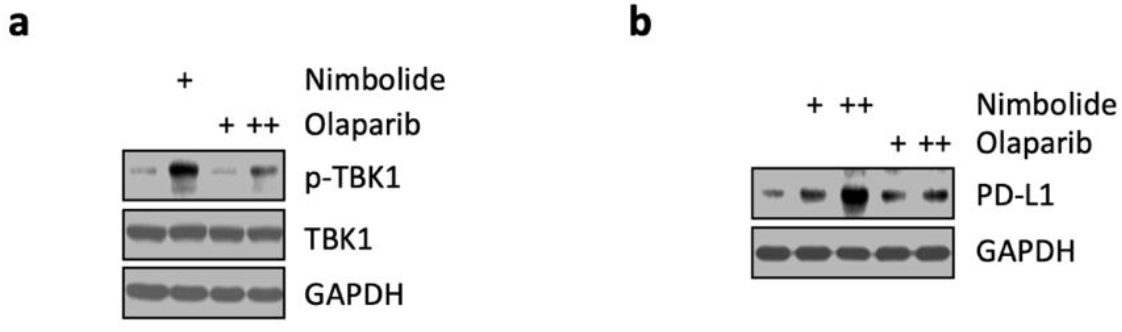
Nimbolide treatment induces the activation of innate immune signaling. **a,** Nimbolide induces strong activation of innate immune signaling. HeLa cells were treated with nimbolide (1 μM, +) or Olaparib (1 μM, + or 10 μM, ++) for 48 h. The whole cell lysates were subject to immunoblot experiments using the indicated antibodies. **b,** nimbolide induces expression of PD-L1. UWB1 cells were treated with nimbolide (1 μM, + or 2 μM, ++) or Olaparib (5 μM, + or 10 μM, ++) for 48 hours. The whole cell lysates were subject to immunoblot experiments using the indicated antibodies.

**Extended Data Fig. 6.**
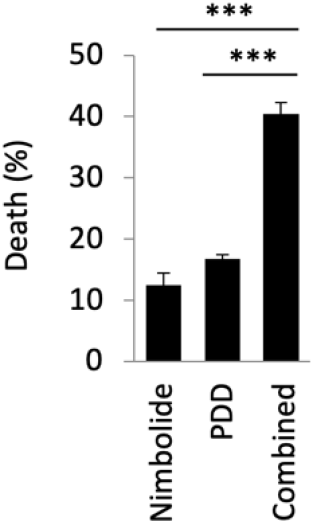
Nimbolide synergizes with a PARG inhibitor. The synergistic effects between nimbolide and PDD00017273 (a PARG inhibitor, PDD). UWB1 cells were treated with either nimbolide (0.25 μM) alone, PDD00017273 (0.25 μM) alone or nimbolide in combination with PDD00017273 (both at 0.25 μM) for 96 hrs.

