## Supplementary information for "Nimbolide Targets RNF114 to Induce the Trapping of PARP1 and Poly-ADP-Ribosylation-Dependent DNA Repair Factors"

**Compound 3**

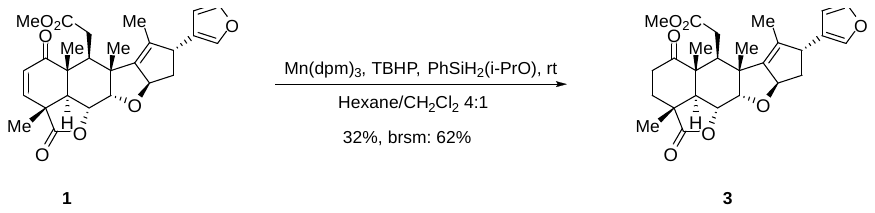

Nimbolide (20 mg, 0.0429 mmol) and Mn(dpm)_3_ (1.3 mg, 0.0021 mmol, 5%) was dissolved in 2 mL hexane and 0.5mL CH_2_Cl_2_ under Ar. PhSiH_2_(OiPr) (15μL, 0.0858 mmol, 2 eq) and TBHP (5.5 M in decane, 16 μL, 0.0858 mmol, 2 eq) were added in sequence. The mixture was stirred at room temperature for 50 min. Saturated Na_2_S_2_O_3_ were added. The layers were separated and the aqueous layer was extracted several times with EtOAc. The combined organic extracts were washed with brine, dried over MgSO_4_, and concentrated. Purification by silica gel chromatography (hexanes:EA = 2:1) afforded the product as white solid 6.4 mg (yield 32%, brsm: 62%) and nimbolide 9.6 mg (yield 48%).

**[α] _D_^25^** = + 119.22 (c 0.253, MeOH), (literature**[α] _D_^25^** = + 122.2 (c 0.1, MeOH)^[1]^)

**^1^H NMR (600 MHz, CDCl_3_):** 7.33 (t, *J* = 1.7 Hz, 1H), 7.26 – 7.24 (m, 1H), 6.32 (dd, *J* = 1.9, 0.9 Hz, 1H), 5.53 (ddt, *J* = 8.4, 6.6, 1.9 Hz, 1H), 4.56 (dd, *J* = 12.1, 3.5 Hz, 1H), 4.21 (d, *J* = 3.5 Hz, 1H), 3.67 (dd, *J* = 8.5, 1.8 Hz, 1H), 3.56 (s, 3H), 2.86 (dd, *J* = 15.7, 5.2 Hz, 1H), 2.81 (ddd, *J* = 16.2, 11.6, 8.5 Hz, 1H), 2.71 – 2.67 (m, 2H), 2.40 – 2.35 (m, 1H), 2.32 (dd, *J* = 15.7, 5.8 Hz, 1H), 2.22 (dd, *J* = 12.1, 6.7 Hz, 1H), 2.14 – 2.08 (m, 3H), 1.70 (d, *J* = 1.8 Hz, 3H), 1.50 (s, 3H), 1.33 (s, 3H), 1.28 (s, 3H) ppm.

**^13^C NMR (150 MHz, CDCl_3_):** δ 210.42, 177.70, 172.90, 144.99, 143.01, 138.89, 135.97, 126.52, 110.38, 88.31, 82.77, 72.71, 51.64, 50.02, 49.60, 49.48, 49.33, 41.16, 40.79 (2C), 34.37, 33.22, 32.87, 17.06, 15.72, 15.09, 12.81 ppm.

HRMS (ESI-TOF): calc’d for C_27_H_32_O_7_ [M+H]^+^: 469.2216, found: 469.2221.

**TLC:** R*_f_* = 0.4 (1:1 hexanes : ethyl acetate).

**Compound 2**

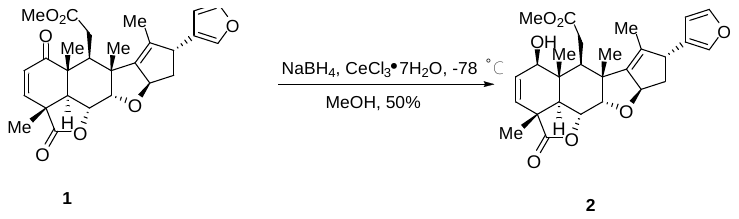

Nimbolide (25 mg, 0.054 mmol) was dissolved in 3 mL MeOH. The mixture was cooled to -78 ℃ and CeCl_3_·7H_2_O (40.3 mg, 0.108 mmol, 2 eq) was added followed by NaBH_4_(4.9 mg, 0.108 mmol, 2 eq). After stirring at -78 ℃ for 30 min, the reaction was quenched by 30 ml saturated NH_4_Cl then warm to rt another 30 ml H_2_O was added. The layers were separated and the aqueous layer was extracted with EtOAc (30 mL × 6). The combined organic extracts were washed with brine, dried over MgSO_4_, and concentrated give the crude. Purification by silica gel chromatography (hexanes:EA 1:1.5) afforded the product as white soild 12 mg (yield 50%).

**[α] _D_^26^** = + 38.01 (c 0.1, CHCl_3_).

**^1^H NMR (400 MHz, CDCl_3_):** δ 7.34 (t, *J* = 1.7 Hz, 1H), 7.21 (s, 1H), 6.24 (d, *J* = 1.6 Hz, 1H), 6.16 (dd, *J* = 9.9, 2.4 Hz, 1H), 5.50 (dd, *J* = 9.9, 2.4 Hz, 1H), 5.48 – 5.41 (m, 1H), 4.51 (dd, *J* = 12.3, 3.7 Hz, 1H), 4.24 (dd, *J* = 8.0, 3.1 Hz, 2H), 3.67 (d, *J* = 8.6 Hz, 1H), 3.54 (s, 3H), 2.82 (dd, *J* = 15.5, 6.3 Hz, 1H), 2.56 (d, *J* = 12.3 Hz, 1H), 2.40 (dd, *J* = 15.5, 5.2 Hz, 1H), 2.30 (t, *J* = 5.7 Hz, 1H), 2.23 (dd, *J* = 12.1, 6.6 Hz, 1H), 2.17 – 2.10 (m, 1H), 1.74 (d, *J* = 1.9 Hz, 3H), 1.35 (s, 3H), 1.32 (s, 3H), 1.06 (s, 3H) ppm.

**^13^C NMR (100 MHz, CDCl_3_):** δ 176.74, 174.75, 145.54, 143.29, 138.95, 136.16, 133.14, 130.66, 126.84, 110.53, 88.33, 83.06, 74.48, 52.14, 50.29, 49.60, 47.55, 46.73, 43.61, 41.41, 40.75, 32.20, 29.85, 19.15, 16.64, 13.09, 12.81 ppm.

HRMS (ESI-TOF): calc’d for C_27_H_32_O_7_ [M+H]^+^: 469.2221, found: 469.2221.

TLC: R*_f_* = 0.4 (EA:hexanes 1:1)

**Compound 4**

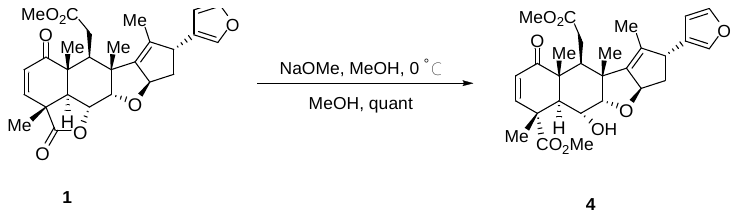

Under Ar (g), the nimbolide (20.8 mg, 0.0446 mmol) and NaOMe (7.2 mg, 0.13 mmol, 3 eq) were dissolved in 3 mL MeOH at 0 ℃ stir at 0 ℃ for 1 hour, the reaction complete, remove the solvent under vacuo give the crude then purified on silica gel chromatography (hexanes:EA 1:1.5) afford the product as white solid 22.1 mg (quant).

**[α] _D_^24^** = +101.16 (c 0.17, CHCl_3_) (literature^[2]^ **[α] _D_^20^** = + 110 (c 1, CHCl_3_)

**^1^H NMR (600 MHz, CDCl_3_):** δ 7.33 (t, *J* = 1.7 Hz, 1H), 7.24 (d, *J* = 1.1 Hz, 1H), 6.41 (d, *J* = 10.1 Hz, 1H), 6.33 (dd, *J* = 1.9, 0.9 Hz, 1H), 5.85 (d, *J* = 10.1 Hz, 1H), 5.55 (ddt, *J* = 8.4, 6.6, 2.0 Hz, 1H), 4.02 (d, *J* = 3.3 Hz, 1H), 3.92 (dd, *J* = 11.7, 3.3 Hz, 1H), 3.70 (s, 3H), 3.67 (s, 1H), 3.66 (s, 3H), 3.40 (d, *J* = 11.7 Hz, 1H), 2.90 (dd, *J* = 16.4, 5.7 Hz, 1H), 2.76 (dd, *J* = 5.7, 3.8 Hz, 1H), 2.26 – 2.20 (m, 1H), 2.20 – 2.17 (m, 1H), 2.04 (dt, *J* = 11.9, 8.5 Hz, 1H), 1.68 (d, *J* = 1.9 Hz, 3H), 1.59 (s, 3H), 1.29 (s, 3H), 1.21 (s, 3H) ppm.

**^13^C NMR (150 MHz, CDCl_3_):** δ 202.25, 175.61, 173.73, 148.14, 146.84, 143.12, 139.05, 134.98, 126.86, 126.47, 110.48, 87.44, 86.97, 66.25, 53.08, 51.73, 49.65, 47.79, 47.51, 47.37, 43.68, 41.48, 39.10, 34.42, 17.58, 17.19, 16.45, 12.91ppm.

HRMS (ESI-TOF): calc’d for C_28_H_34_O_8_ [M+H]^+^: 499.2319, found: 499.2326.

TLC: R*_f_* = 0.4 (CH_2_Cl_2_:EA 6:1)

**Compound 5**

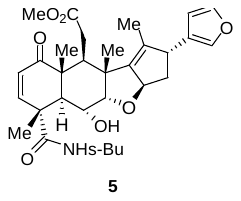

The compound 5 was prepared following reported procedure^[3]^.

^1^H NMR spectra of compound **3** in CDCl_3_ (600 MHz)

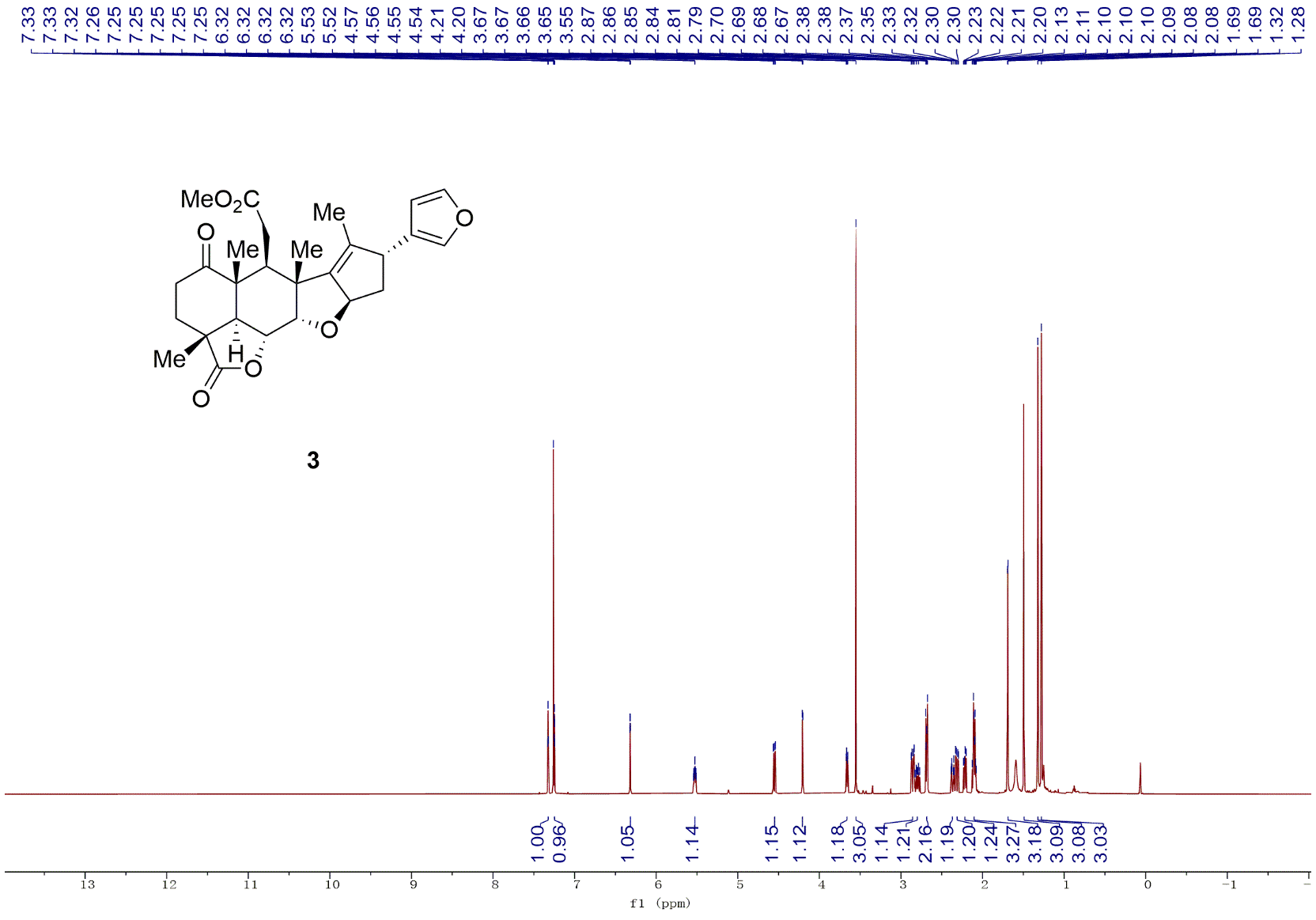

^13^C NMR spectra of compound **3** in CDCl_3_ (150 MHz)

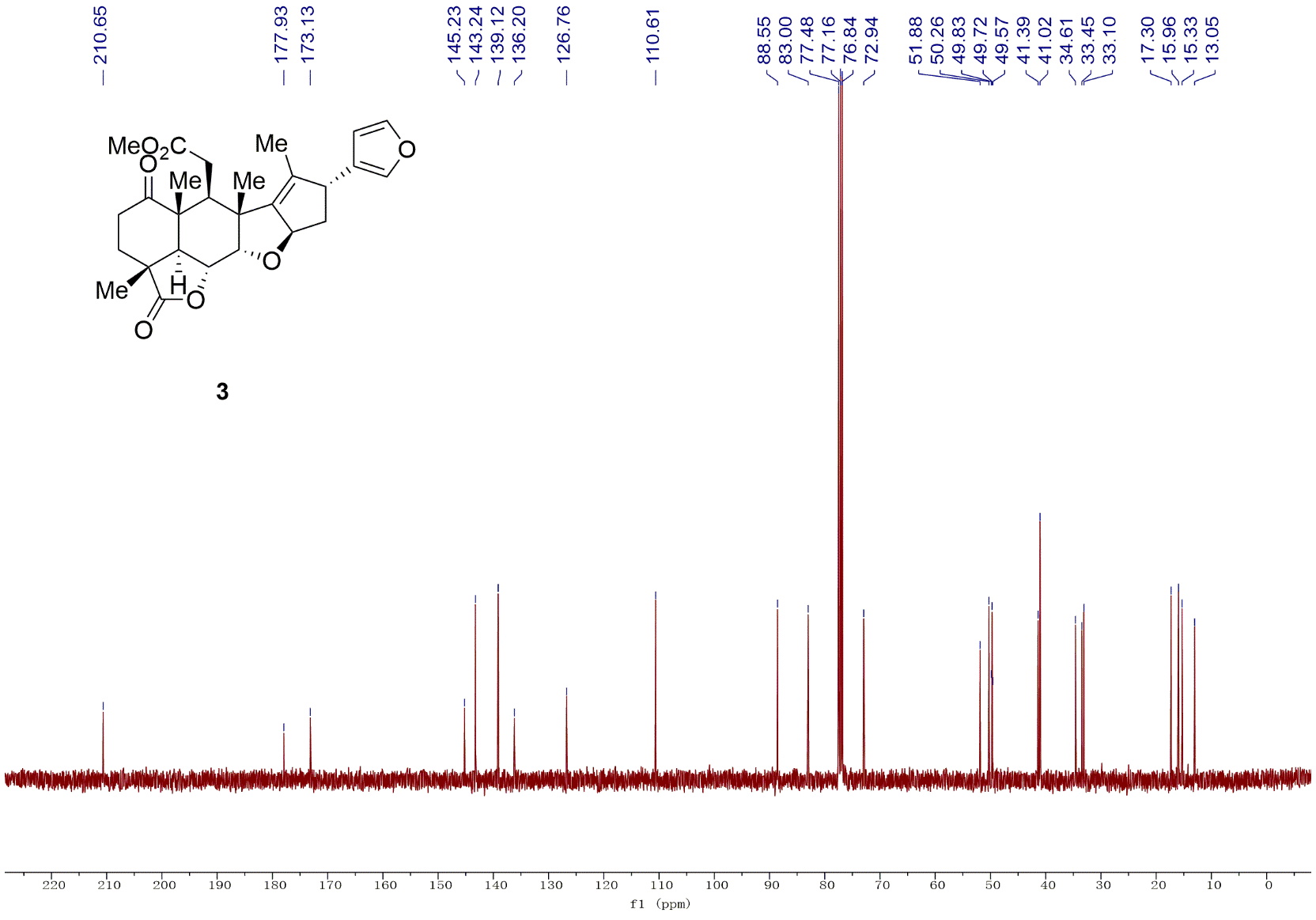

^1^H NMR spectra of compound **2** in CDCl_3_ (400 MHz)

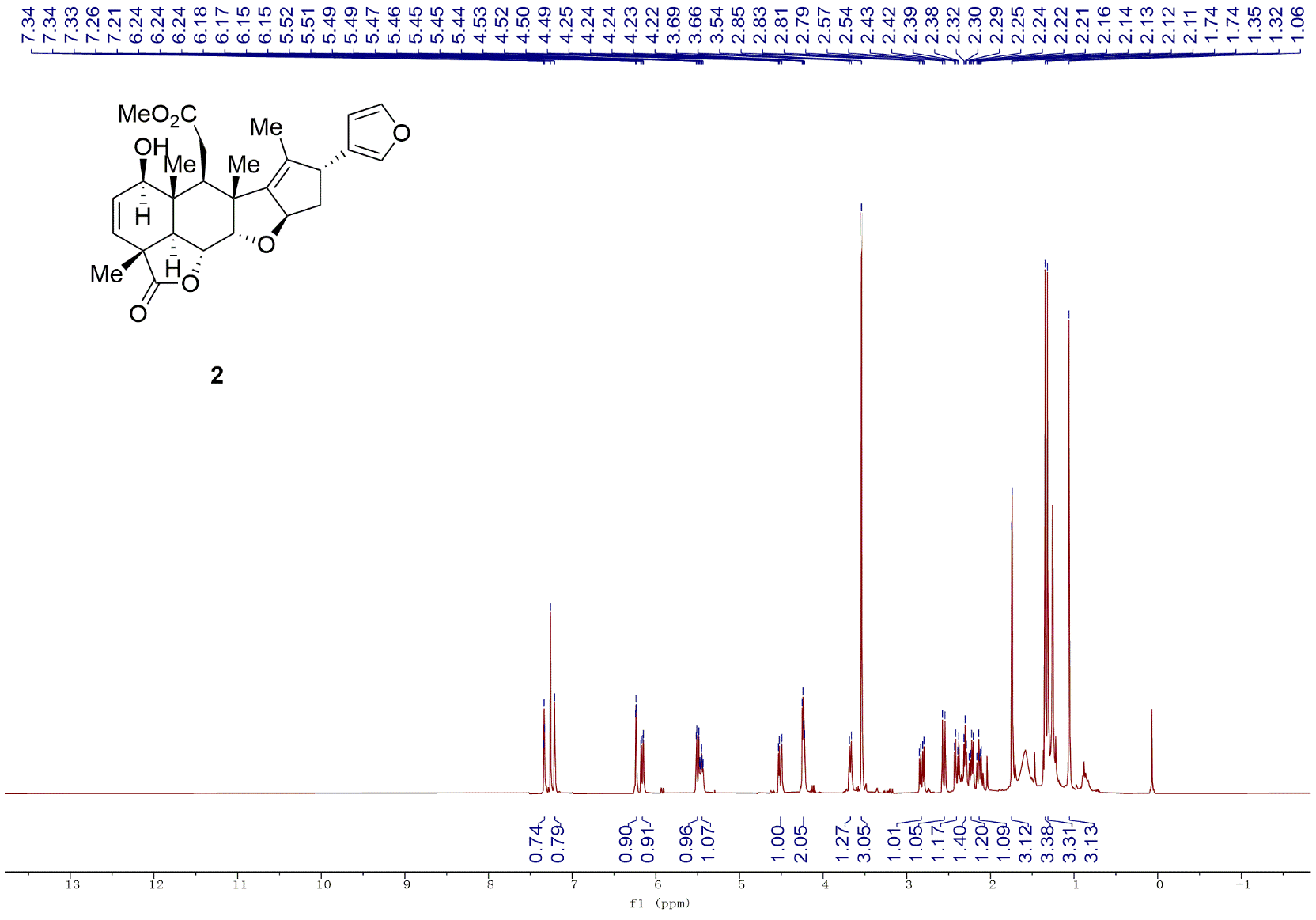

^13^C NMR spectra of compound **2** in CDCl_3_ (100 MHz)

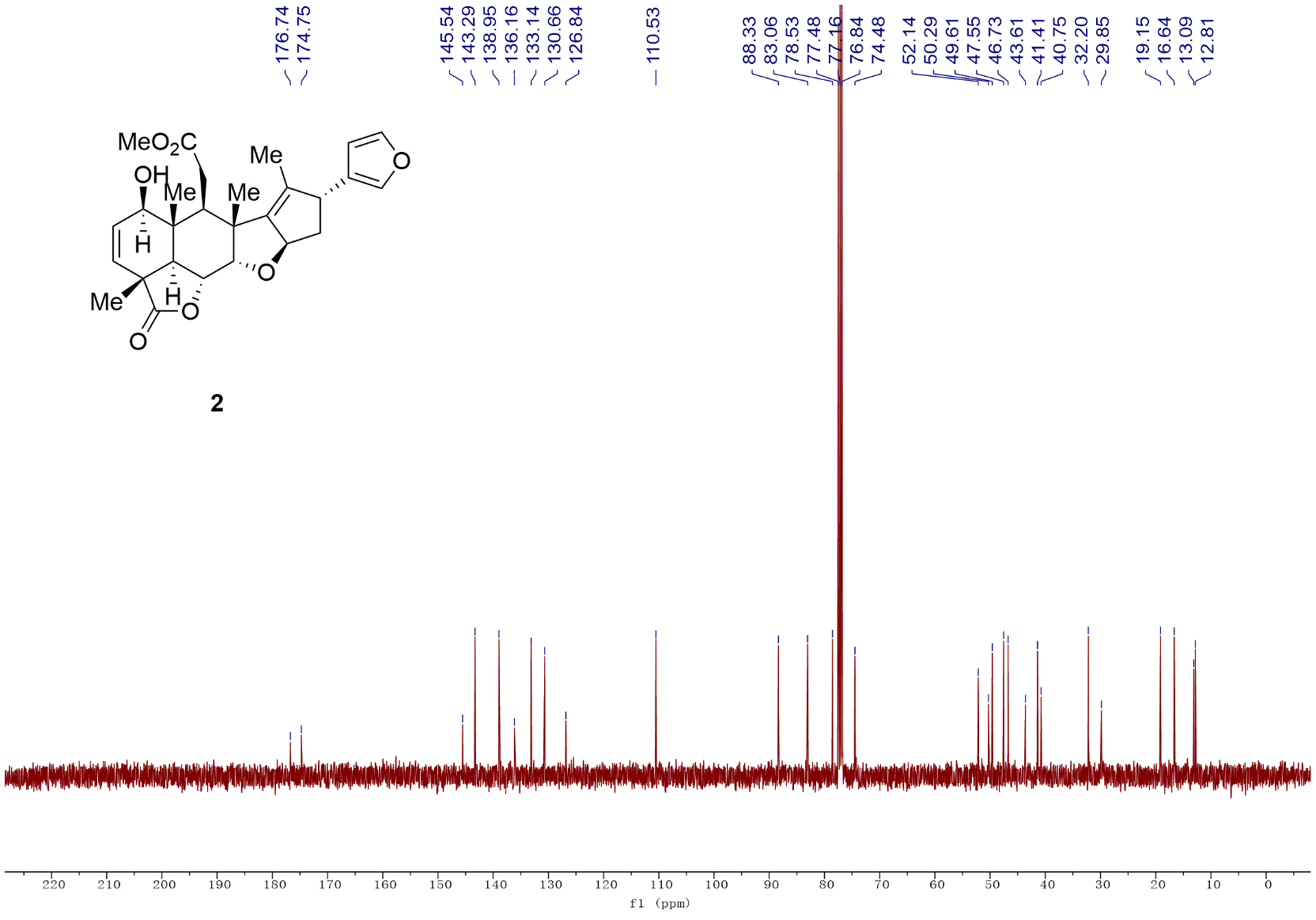

H-H NOESY of compound **2** in CDCl_3_ (400 MHz)

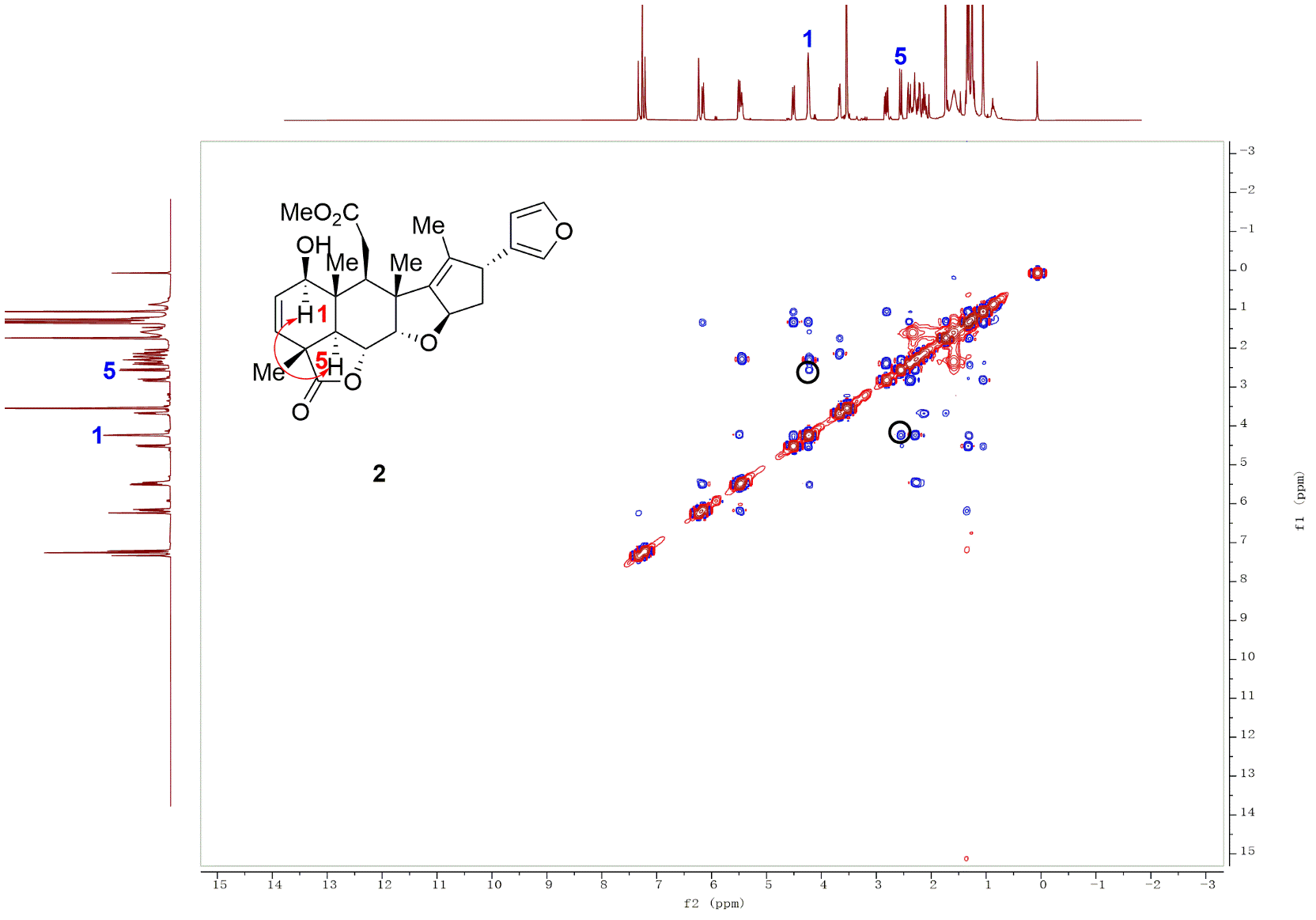

^1^H NMR spectra of compound **4** in CDCl_3_ (600 MHz)

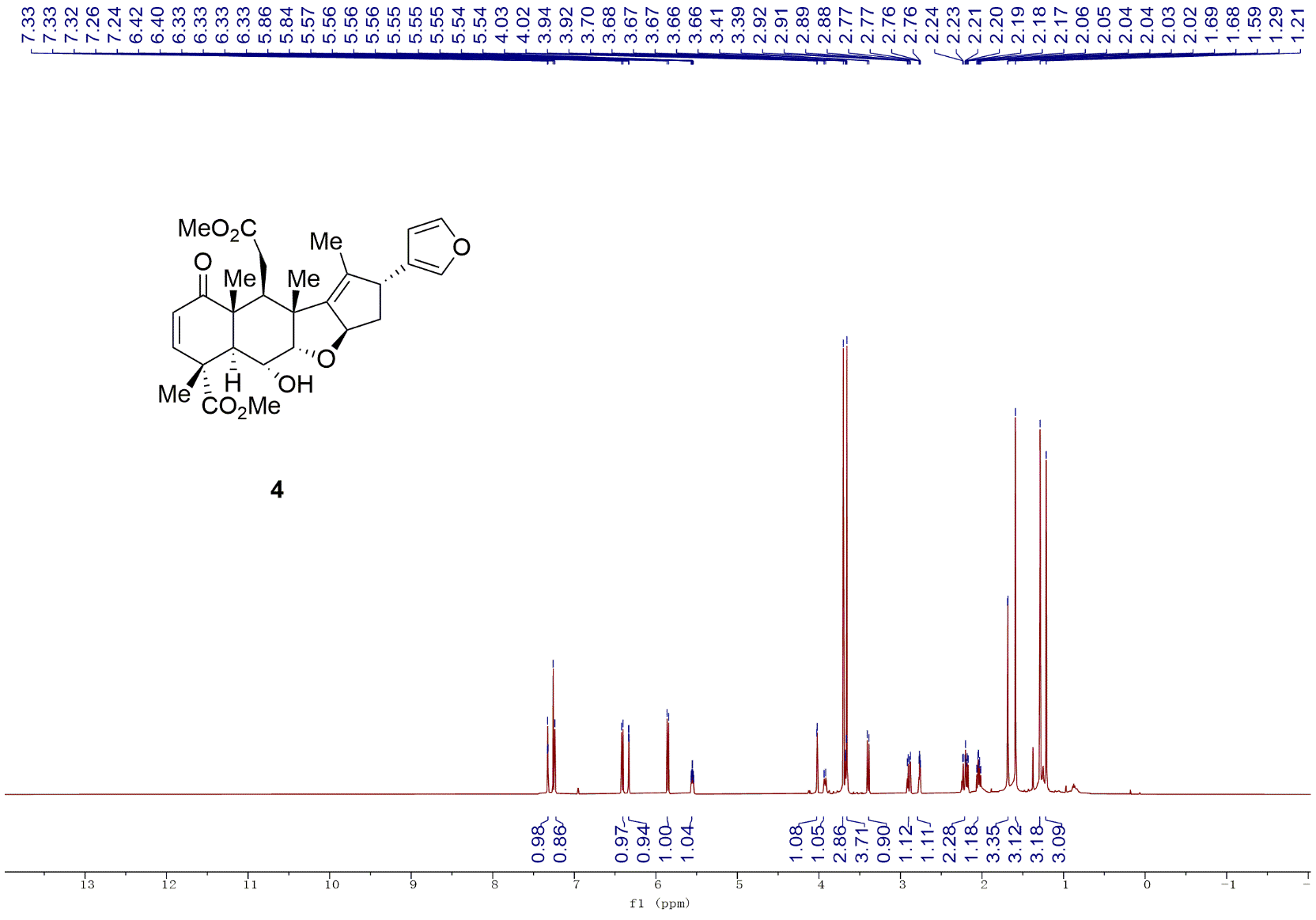

^13^C NMR spectra of compound **4** in CDCl_3_ (150 MHz)

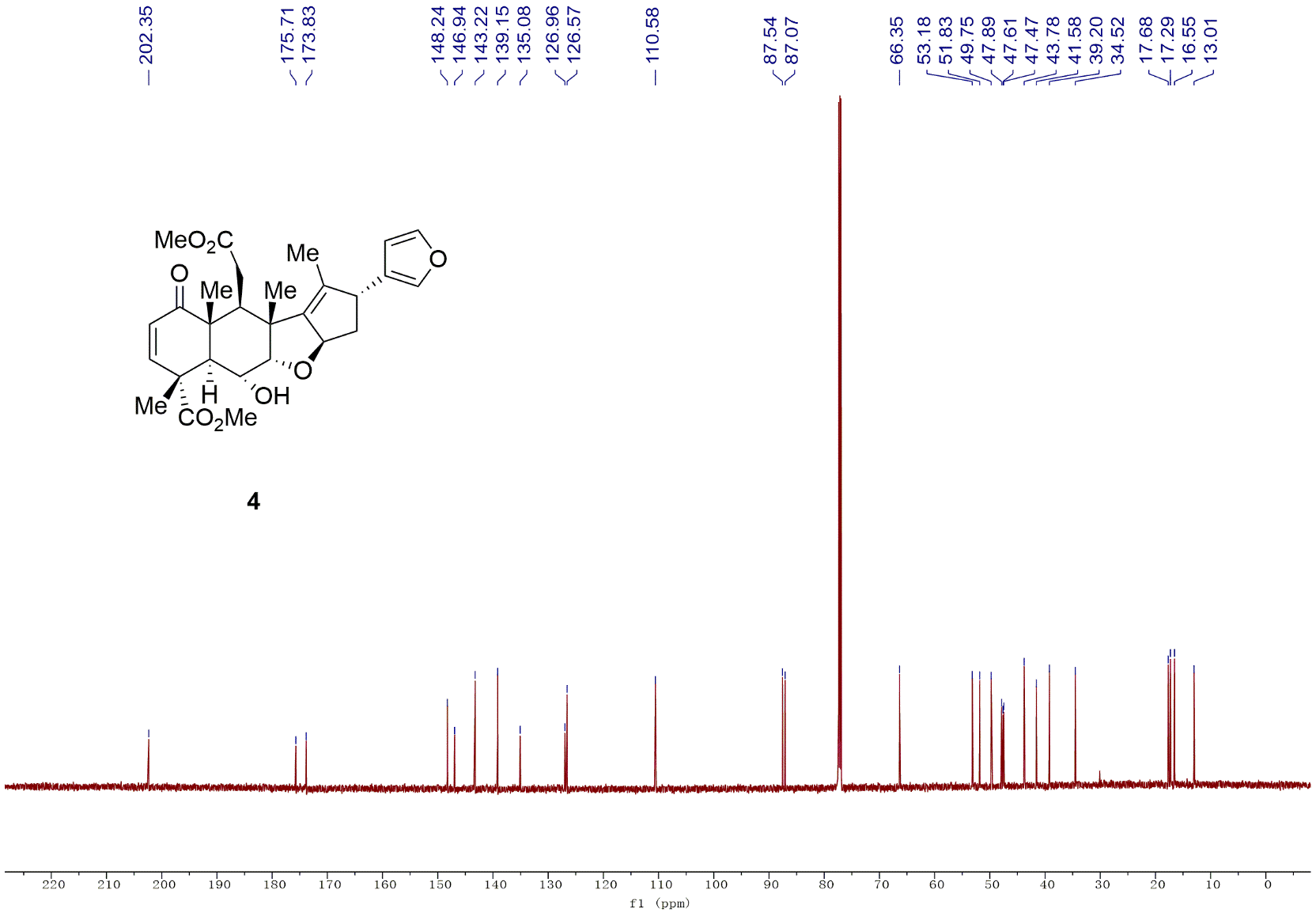
